# Fluctuating environments select for short-term phenotypic variation leading to long-term exploration

**DOI:** 10.1101/394676

**Authors:** Rosangela Canino-Koning, Michael J. Wiser, Charles Ofria

## Abstract

Genetic spaces are often described in terms of fitness landscapes or genotype-to-phenotype maps, where each genetic sequence is associated with phenotypic properties and linked to other genotypes that are a single mutational step away. The positions close to a genotype make up its “mutational landscape” and, in aggregate, determine the short-term evolutionary potential of a population. Populations with wider ranges of phenotypes in their mutational neighborhood are known to be more evolvable. Likewise, those with fewer phenotypic changes available in their local neighborhoods are more mutationally robust. Here, we examine whether forces that change the distribution of phenotypes available by mutation profoundly alter subsequent evolutionary dynamics.

We compare evolved populations of digital organisms that were subject to either static or cyclically-changing environments. For each of these, we examine diversity of the phenotypes that are produced through mutations in order to characterize the local genotype-phenotype map. We demonstrate that environmental change can push populations toward more evolvable mutational landscapes where many alternate phenotypes are available, though purely deleterious mutations remain suppressed. Further, we show that populations in environments with harsh changes switch phenotypes more readily than those in environments with more benign changes. We trace this effect to repeated population bottlenecks in the harsh environments, which result in shorter coalescence times and keep populations in regions of the mutational landscape where the phenotypic shifts in question are more likely to occur. Typically, static environments select solely for immediate optimization, at the expensive of long-term evolvability. In contrast, we show that with changing environments, short-term pressures to deal with immediate challenges can align with long-term pressures to explore a more productive portion of the mutational landscape.

## Introduction

Interactions are ubiquitous in evolving systems. Some of these interactions are between individuals of the same population [1–5]; others are between members of different populations [6–8]. A third group – interactions between an individual and the environment [9–11] – can also be crucial, both for how an individual experiences the world, and how it modifies its surroundings [12, 13], which can have impacts on the rest of the ecosystem [14, 15].

The interactions between an environment and possible genomes can be mathematically expressed by a fitness landscape. Fitness landscapes are a mathematical tool to map genetic sequences to reproductive fitness. Many studies have examined the important role that different types of fitness landscapes play on evolutionary dynamics and outcomes, both in biological populations [16–19] and in evolutionary computation settings [20–22]. However, real-world fitness landscapes are far more complex and varied than the limited or idealized models that are used in most of these studies. Neighboring regions of real landscapes can have starkly different properties from each other based on the effects of and interactions among mutations; as such, a local region of a fitness landscape around a genotype is commonly referred to as its mutational landscape.

Different landscapes, or different regions of a landscape, can vary tremendously in their properties. Examples of the type of properties that we are interested in include robustness, epistasis, and modularity, all of which are measurements of how information is organized inside of a genome and commonly categorized as components of an organism’s “genetic architecture”. Isolated pockets in a landscape can often be characteristically different from the landscape as a whole due to the amount and organization of genetic information. In fact, in most natural fitness landscapes, the vast majority of neighborhoods consist entirely of non-replicating genomes with zero fitness (and thus no genetic information), making life itself appear to be a rare exception [23].

Evolution on realistic landscapes is clearly limited to those regions that have non-zero fitness, with a selective pressure for fitness to increase. Beyond evolution away from zero-fitness regions, populations can evolve in more complicated ways, toward neighborhoods with specific local properties based on the evolutionary forces acting upon the populations. For example, high mutation rates drive populations toward neighborhoods with a higher fraction of neutral mutations in an effect dubbed “survival of the flattest” [24]. Similarly, sexual populations tend toward regions of the fitness landscape with more modularity [25] and more negative epistasis [26] than otherwise equivalent asexual populations.

Understanding the dynamics of evolution in complex meta-environments, such as changing environments, is of broad interest. It is important to evolutionary computation, given the strong influence of local landscape properties on the quality of the final solutions that an evolving population is able to obtain. Its relevance to evolutionary biology is equally obvious – the local landscape that a population occupies will influence the selective forces at play in the population, creating a feedback cycle between these two important evolutionary factors [8, 27–29]. Disentangling such interactions is likely to provide further insights into fundamental evolutionary dynamics. Computational artificial life systems have the advantage of being able to bridge these two realms: they have unconstrained evolutionary dynamics similar to natural systems, while maintaining the ability to rapidly perform experiments and collect any data we need about populations or their local landscapes.

### Evolvability and Genetic Architecture

Evolvability refers to a series of distinct but overlapping concepts that are generally concerned with adaptation, variation, and/or novelty generation [30]. Depending on your perspective, evolvability can describe the response to selection at the population level [31, 32], the ability of populations to adapt to changing conditions [33], larger phenomena such as variability generation [34], exploration of neutral spaces and robustness [35, 36], generation of novel features [37, 38], or even the potential to generate clade-level innovations [39] and major transitions [40]. Here, we will focus on evolvability as the capacity for mutations to generate adaptive variation in a genome.

In the short-term, this kind of evolvability determines a population’s response to selection, and depends primarily on the organization and interrelation of information in the genome; that is, the genetic architecture, and the resulting genotype-to-phenotype map [34]. An example of evolvable architecture can be found in some bacterial genomes that contain highly mutable genome regions, called contingency loci. Small sets of insertions or deletions to these regions create transcription frameshifts that alter the expression of nearby coding regions, thus allowing populations to easily switch phenotypes via minor mutations. Contingency loci are most often seen in the genomes of pathogens, which are subject to frequent environmental shifts caused by the host immune system [41]. Thus, these populations are able to produce large amounts of heritable variation despite their reduction in diversity resulting from population bottlenecks.

### Mutational Landscapes

Properties of genetic architectures such as evolvability and robustness are determined by the shape of the resulting mutational landscape (local fitness landscape around a genotype, accessible in a single mutation) [42]. Robust genetic architectures that can tolerate more mutations without altering their phenotype reside in mutational landscapes that connect to more neutral mutants. Similarly, architectures where mutations are more likely to cause phenotype switching without substantial reductions in fitness, reside in more evolvable regions of genotype-space.

It is worth noting that not all neighborhoods of the mutational landscape may be equally accessible. Some genome regions may be more robust to mutation than others. For example, in *E. coli*, the methyl-directed mismatch repair (MMR) pathway has been shown to preferentially repair coding regions over non-coding regions [43]. Alternately, some kinds of mutations may be more likely to occur than others. A mutation accumulation (MA) study of *S. typhimurium* found a strong bias toward GC-to-TA transversions rather than GC-to-AT transitions [44]. These kinds of effects thereby skew the probabilities of some kinds mutations occurring that might lead into certain neighborhoods of the mutational landscape. These kinds of differential probabilities may therefore moderate a population’s diffusion through the mutational landscape.

Further, response to selection is likely to be weaker in regions of the landscape where there are fewer available mutations that provide potentially adaptive traits, whereas response to selection will be stronger in regions where there are many adaptive variants available within a few mutational steps [37, 45]. This differential response to selection may therefore constrain the ability of populations to diffuse across a fitness landscape.

### Landscape Metrics

Assessing the qualities of the nearby mutational landscape requires measures that can relate phenotypes and their fitness effects with the probabilities that these mutants will arise in the population. For the purposes of this paper, we define the organism phenotype as being the set of logical tasks performed by an organism. Phenotype contributes to fitness, but fitness is a distinctly calculated value. In order to assess the relative neutrality of the nearby mutational network, we will measure the **Genomic Diffusion Rate** *D*_*g*_ [46]. This rate approximates the overall rate at which the population encounters new neutral genotypes. To measure the **Genomic Diffusion Rate** (*D*_*g*_) in the local neighborhood of a genotype, we first calculated its *Fidelity* (*F*), or the probability of an offspring sharing this genotype with its parent. Given a uniform mutation rate, Fidelity is the probability that a single locus is not mutated, (1 *-µ*), raised to the power of the genome length (*l*)^1^.

Next, we measured the proportion of one-step mutants that were neutral or beneficial when compared to the parent (*p*_*v*_) as well as those that were detrimental or lethal (*p*_*d*_), which must sum to one (*p*_*v*_ + *p*_*d*_ = 1). The *Neutral Fidelity* (*F*_*v*_) of a genotype is thus the probability that no harmful mutations occur, assuming no epistasis. Finally, subtracting Fidelity from Neutral Fidelity yields the overall probability of producing an offspring with a different genotype, yet neutral or better fitness (*D*_*g*_).

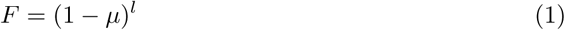

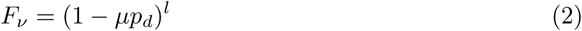

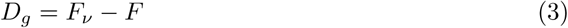

Measures of neutral exploration, however, only show part of the picture. While some form of neutrality is necessary for exploring a fitness landscape, new phenotypes must be discoverable to achieve higher local evolvability. In order to assess evolvability more specifically, we introduce a related measure, the **Phenotypic Diffusion Rate** (*D*_*p*_), which represents the probability that an offspring will be fitness-neutral (or better), but also express a different phenotype than its parent. To do so, we must first measure the proportion of one-step mutants that are *phenotypically* neutral as compared to their parent (*p*_*pv*_) and follow a similar procedure as above, first calculating the probability that a phenotype-changing mutation will occur (*µ*_*pheno*_), then the phenotypic-level Fidelity (*F*_*pv*_).

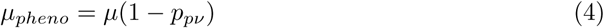

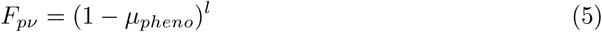

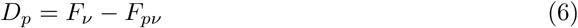

The difference between the overall Neutral Fidelity and the phenotype-preserving Neutral Fidelity (*F*_*v*_ - *F*_*pv*_) yields the phenotypic diffusion rate.

### Expected Value of Fitness Landscapes

In the context of changing environments, the expected fitness value (*E*(*w*)), and thus the neutrality, of a mutant on the mutational landscape will vary depending on the environmental context. So, in one environment, a mutant may be highly fit, but the same allele may be highly deleterious in a different environment. In order to address this variation, all metrics must be normalized by the probability that a particular environment will occur (*P*_*i*_). That is, the nearby mutational landscape must be evaluated in each possible environment, yielding a traditional fitness landscape. Then, the set of fitnesses of each mutant (*w*_*i*_) in each environment must be aggregated according to the probability of that environment occurring.

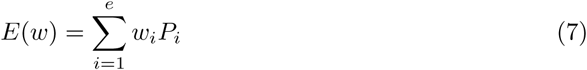

### Changing environments create more paths to different kinds of phenotypes

Sustained directional selection adjusts the composition of phenotypes and genotypes in a population [47], typically moving that population across the mutational landscape to local regions of higher fitness. When populations find a fitness peak, they tend to cluster there, and exploration of regions further away slows dramatically.

In changing environments, however, the direction of selection is not fixed and peaks are not stable. Instead, as the environment changes, populations are driven to explore new regions of the mutational landscape [48–50]. As they proceed, populations accumulate and carry with them the genetic material acquired in prior explorations and adaptations, and use this history as raw material for new adaptation [51]. Indeed, earlier work has shown that changing environments promote evolvability in many contexts [48], without compromising robustness [24, 52]. Strength of selection is also an important component of this exploration, since the harshness of the environment drives the speed with which organisms adapt to new conditions [53].

For longer evolutionary timescales, beyond the limited scope of direct response to selection against an environment, evolvability is concerned with generation of variability and exploration of neutral spaces. Populations that exhibit this kind of evolvability would possess genomes with genetic architectures that more easily traverse the mutational landscape along neutral roads and thereby discover new fitness peaks while avoiding needing to cross fitness valleys. This kind of evolvability would allow populations to more easily colonize new ecological niches and form new clades [38, 39].

Despite some common features, the relationship between short-term and long-term evolvability is not obvious. Architectural features and evolutionary pressures that convey short-term evolvability may not be the same as those that confer longer-term evolvability [30]. For example, features such as anti-robustness that promote rapid adaptation to a harsh fluctuating environment might reduce fitness in constant or benign fluctuating environments as compared to that of wild-type invaders. Alternately, the adaptation to harsh fluctuating environments and the resulting bottlenecks would potentially reduce diversity to the point where large amounts of neutral novelty generation could not occur.

Finally, there is some evidence that the types of selection regimes typically used in experiments with changing environments and evolvability might preferentially favor individual evolvability (the probability of an individual’s offspring accessing novel phenotypes) over population-level evolvability (the probability of the population at large accessing novel phenotypes) [54, 55]. Adaptive selection – that is, selection toward a particular goal – has been shown to depress population diversity even while it increases individual evolvability in changing environment regimes. In contrast, divergent (diversity-promoting) selection, such as frequency-dependent selection, increases standing diversity, and thus evolvability at the population level [54]. Therefore, it is not clear that the kinds of selective pressures that promote short-term adaptation in changing environments would, in turn, promote exploration and exploitation of novel environments.

In this paper, we show how changing environments not only drive exploration of the mutational landscape, but also select for populations whose genetic architectures are qualitatively different than those from populations evolved in static environmental conditions under purely directional selection.

In particular, we show that populations evolved under harsh, cyclically-changing environments have many more changes along their phylogenetic histories than those evolved in static or benign changing environments. Organisms evolved in these populations also contain reservoirs of pseudogene-like vestigial loci that were acquired and deactivated through repeated adaptation and fixation cycles. As a result, populations evolved in these harsh cyclically-changing environments are low in standing neutral diversity at the population level, but they still connect locally with many more phenotypically-interesting regions of the mutational landscape than more diverse populations evolved in static or benign environments.

Even so, we show that the strong selective pressures associated with these harsh environments are detrimental to long-term evolvability, and instead, that benign environments, with their higher standing diversity, are more successful at adapting to entirely new environments.

### Digital Evolution

Digital Evolution uses self-replicating computer programs as model organisms to study evolutionary dynamics [56]. Unlike theoretical simulations, digital organisms have a fully functional genome that direct them to self-replicate, mutate, and compete with their peers for resources and space in which to reproduce. Because digital organisms undergo random genetic mutations (i.e., variation) that are passed on to their offspring (inheritance), and their survival is based on the actions they take (differential selection), they undergo evolution by natural selection [57].

Indeed, because evolution is an algorithmic process, studies with digital organisms are not simulations of evolution, but actual instantiations of evolution, albeit on an artificial substrate. Therefore, research performed using digital organisms are not theoretical explorations, but rather true experiments, where hypotheses are tested, and the outcomes are not pre-arranged. Studies of evolutionary processes in digital organisms are particularly well suited to examining fundamental questions about evolutionary principles, such as how information flows through evolutionary processes, how arrangements of genetic architecture can affect evolutionary trajectories, how different types of selective pressures interact to produce complexity, to name a few.

Digital organisms do not suffer from many of the drawbacks of experimentation on natural organisms. Three of the advantages of digital organisms are particularly relevant for our study. First, the rates of reproduction in digital systems are much faster than in even the most rapidly-reproducing physical organisms; we can process generations of organisms in seconds, rather than the hours required for the fastest biological organisms under sustained conditions [58, 59], or the weeks to years needed for more complex multicellular organisms [60, 61].

Second, using digital organisms allows us to tightly control and verify experimental conditions. For example, in physical organisms, factors such as mutation rate can generally be measured only after the fact, or coarsely altered through mutagens. In digital organisms, however, we can not only control mutation rates with fine-grained precision, but also types and probabilities of different types mutations (e.g., substitutions vs. insertions vs. deletions). Furthermore, we are also able to track and replay the evolutionary history of every organism at any point in time to verify that unusual or unexpected results do not represent measurement error. This ability to exactly replicate evolutionary results at an individual organism level is firmly out of reach for experiments with physical organisms.

Finally, we can precisely and perfectly map the mutational landscape around the genome of a digital organism, and identify the role of every site in its genome [46]; such exhaustive techniques are not feasible in even the simplest physical organisms. All of these factors make digital organisms ideal for studying the effects of changing environments on the mutational landscape.

## Methods

### Avida Digital Evolution Platform

We used Avida [62] to examine the effects of cyclic changing environments on the genomes of evolved digital organisms. Avida is a software platform for performing evolution experiments with digital organisms in a virtual world.

An Avida organism is composed of a circular genome of assembly-like computer instructions that are executed in a virtual CPU (Fig 1). Populations of these organisms are placed in a toroidal world in individual cells where they are allowed to execute, reproduce, compete for space, mutate, and evolve.

**Fig 1.**
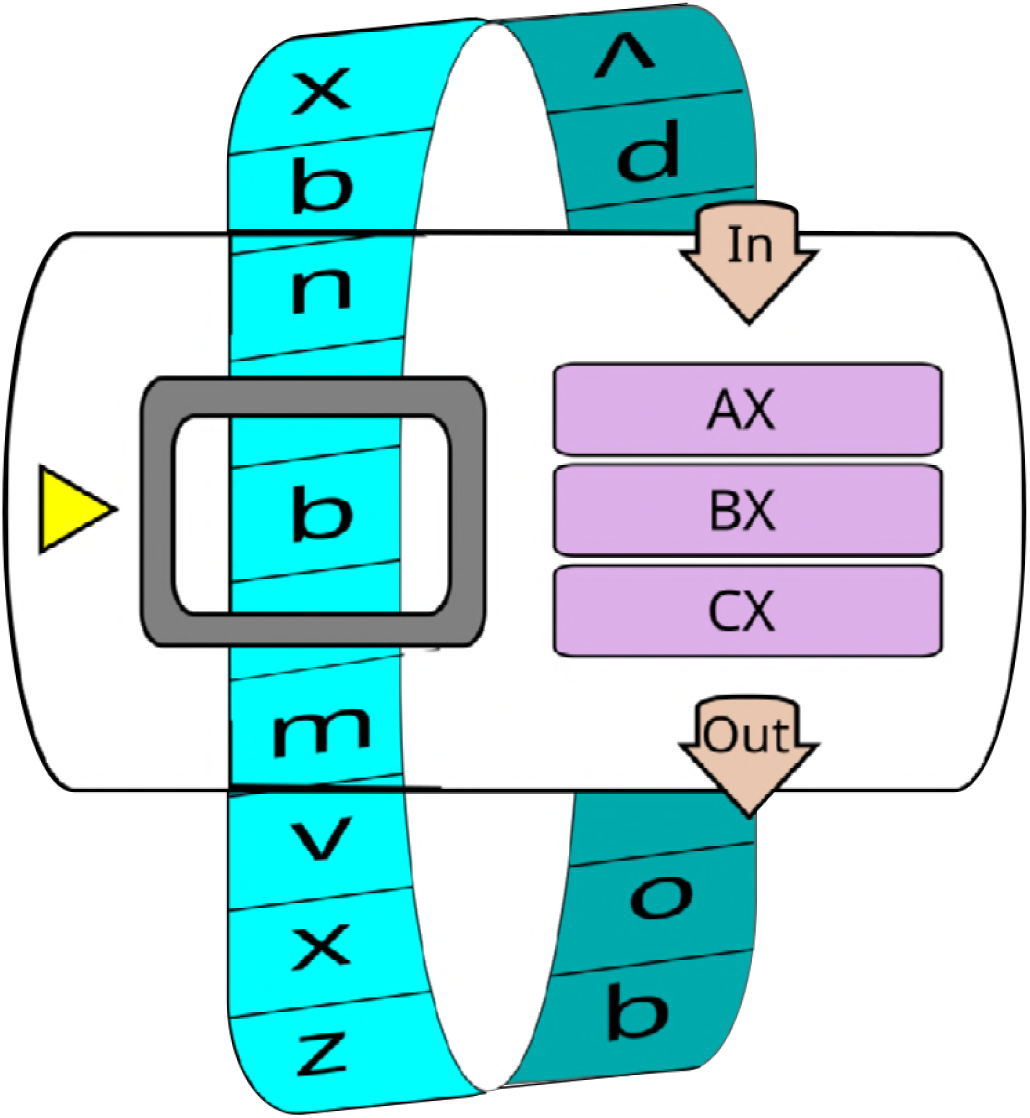
An example virtual CPU from Avida, with a circular genome (blue), three registers (purple), input and output handlers (tan), and an instruction pointer (yellow) indicating the next instruction to be executed [63].

Organisms in Avida are self-replicating, and experience mutation. The genome in the initial default organism contains all of the instructions necessary for reproduction. However, the instructions are not copied into an offspring perfectly. By default, the reproductive copy instruction is faulty, meaning that it will probabilistically introduce errors (mutations) into the offspring genomes. These offspring organisms execute their own genomes even when different from their parent, and in turn pass on their inherited mutations, along with new mutations, to their own offspring (i.e., variation in the systems is heritable).

Avida worlds can be space- or resource-constrained. Avida allows the experimenter to configure many aspects of the environment, thus subjecting the organisms to various kinds of selective pressures. In many cases, these environments will include resources that can be metabolized by performing specific functions or activities, resulting in a boost to execution speed that gives the organisms a competitive advantage. However, even without explicit external pressures, organisms still experience an implicit pressure to execute more quickly and efficiently. The organisms that run fastest are typically able to also reproduce fastest, and thus out-compete their peers for space.

Avida is available for download without cost from http://avida.devosoft.org/, and specific versions along with data-files and analysis scripts to reproduce the experiments described in this paper may be found at https://github.com/voidptr/avida and https://github.com/voidptr/ce_rapid_adaptation_data.

### Experimental Design

In order to examine the dynamics and mechanisms of evolving populations in changing environments, we performed a set of experiments divided into two stages. In the first stage, we subjected populations of evolving digital organisms to a set of benign and harsh cyclic changing environments. The cyclic environments were designed to simulate predictable cycles of change, such as day/night or seasonal cycles. These experiments allow organisms to adapt to a predictable set of environments, and explores short-term evolvability dynamics. See Table 1

**Table 1.** Experimental Treatments - Stage 1 - Cyclic Changing Environments.

| Treatment | Changing Environment | Rewarded Tasks |  |
| --- | --- | --- | --- |
|  |  | XOR | EQU |
| Control | None (static) | constant $2^3$ | constant $2^5$ |
| Benign | Cyclic | constant $2^3$ | benign fluctuating $2^0$ or $2^5$ |
| Harsh | Cyclic | constant $2^3$ | harsh fluctuating $2^{-5}$ or $2^5$ |

Two types of changing environment, plus a static control. In the first two treatments, the environment switches in a predictable cycle. The benign treatment enables and disables reward for the EQU task, while the harsh treatment rewards and then punishes this task.

The second stage takes these change-evolved populations and introduces them to a completely new environment. This set of experiments explores the relationship between evolvability traits acquired via selection for short-term adaptation to cyclical change, and examines how these traits perform in a long-term evolutionary context. See Table 2.

**Table 2.** Experimental Treatments - Stage 2 - Long Term Evolution.

| Treatment | Changing Env. Type | Rewarded Tasks |  |  |  |  |
| --- | --- | --- | --- | --- | --- | --- |
|  |  | Stage 1<br>(0-200,000 Updates) |  | Stage 2<br>(200,000-400,000 Updates) |  |  |
|  |  | XOR | EQU | XOR | EQU | Expanded Task-set<br>(Logic-77 minus XOR & EQU) |
| Control | None (static) | constant $2^3$ | constant $2^5$ | constant $2^3$ | constant $2^5$ | constant $2^{0.3}$ |
| Benign | Cyclic | constant $2^3$ | benign fluctuating $2^0$ or $2^5$ | constant $2^3$ | benign fluctuating $2^0$ or $2^5$ | constant $2^{0.3}$ |
| Benign Quiescent | Cyclic | constant $2^3$ | benign fluctuating $2^0$ or $2^5$ | constant $2^3$ | constant $2^5$ | constant $2^{0.3}$ |
| Harsh | Cyclic | constant $2^3$ | harsh fluctuating $2^{-5}$ or $2^5$ | constant $2^3$ | harsh fluctuating $2^{-5}$ or $2^5$ | constant $2^{0.3}$ |
| Harsh Quiescent | Cyclic | constant $2^3$ | harsh fluctuating $2^{-5}$ or $2^5$ | constant $2^3$ | constant $2^5$ | constant $2^{0.3}$ |

### Short-Term Evolvability - Stage 1

We subjected a total of 150 replicate populations of digital organisms to two different treatments of two-phase cyclically changing environments, plus a static control. The environment cycles between equal-length periods of reward and punishment. Each cycle extends for 1000 updates, or roughly 30 generations. In the static control, there is no cycle. Rather, the rewards remain constant. This stage of the experiment extends for 200 cycles, or 200,000 updates, approximately 6,000 generations.

We set up the system to detect organisms that performed `XOR` or `EQU`, two challenging bit-wise logical tasks. In the static control, `XOR` is rewarded with a CPU speed (and thus fitness) multiple of 8, while `EQU` is rewarded with a CPU speed multiple of 32. In the harsh treatment, as the cycle progresses, the `XOR` reward remains constant, while the `EQU` reward cycles between a 32-fold bonus and a correspondingly harsh 32-fold penalty (i.e., CPU speed is divided by 32 when `EQU` is performed in the off phase of the cycle). The benign treatment is nearly identical to the harsh treatment, except that the reward merely goes away in the off-cycle as opposed to incurring a severe penalty.

In both environments, we identify `EQU` as the *Fluctuating Task*. `XOR`, because it is rewarded continuously, is the *Backbone Task*, and is used as a background for comparing the separation or intertwining of functional genetic components in the evolution of `EQU`. Further, the 4-fold difference in reward level between `XOR` and `EQU` encourages the evolution and maintenance of `EQU` when possible.

### Long-Term Evolvability - Stage 2

The second stage of the experiment continues the evolution of these populations, but introduces them to a completely new environment, with an expanded set of rewarded bitwise tasks to perform: Logic-77 (Table 2). We refer to those tasks which were selected for in stage 1 as the **basic task set**. The Logic-77 task set is a super-set of the basic task set, and includes all bitwise tasks for which there are up to 3 inputs, including those that were initially rewarded in stage 1. We refer to the additional tasks from Logic-77 - those which are not part of the basic tasks set, and that we reward only in stage 2 - as the **expanded task set**. The total Logic-77 task set is a combination of both the basic and expanded task sets.

These new tasks use up to three bit-wise inputs rather than two, and are each rewarded with a constant 1.2-fold bonus to execution. This reward provides a mild selective pressure to evolve these tasks, but the benefits to performing them do not overwhelm the existing selective pressure to continue performing `XOR` or `EQU`.

In order to differentiate between the effects of architectural features and direct effects of alternating selection, we duplicated the populations in each the benign and harsh treatments at the end of stage 1 into two treatments each. Each treatment introduces the rewards of the **expanded task set**, but one treatment in each pair continues the changing environment of the first stage, while the other treatment stops the cycle, and instead rewards the **basic task set** at a constant rate.

- **Static (Control)**: This treatment is a baseline for comparing adaptation to the **expanded task set**. For the first 200k updates (stage 1), we reward populations for performing `XOR` and `EQU` (Table 2). For the second 200k updates (stage 2), we add constant rewards for the **expanded task set**. Each new task is rewarded at a 1.2-fold bonus to task execution.
- **Benign Changing Environment**: This treatment shows the effects of a continuing benign changing environment on adaptation to the **expanded task set**. For the first 200k updates (stage 1), we alternate rewarding and not rewarding populations for performing the `EQU` task (Table 2). For the second stage of the experiment, starting at 200k updates, we add constant rewards for each of the new tasks in the **expanded task set**, at a 1.2-fold bonus to task execution. The environmental fluctuation from the first stage continues through the end of the experiment.
- **Benign Quiescent Changing Environment**: In contrast to the benign changing environment treatment, this treatment tests the abilities of populations initially evolved in a benign changing environment to adapt to the **expanded task set**, but without active environmental fluctuation during the adaptation. For the first 200k updates (stage 1), we alternate rewarding and not rewarding populations, as in the Benign Changing Environment above. For the second stage of the experiment, starting at 200k updates, we add constant rewards for each of the new tasks in the **expanded task set**, at a 1.2-fold bonus to task execution. The environmental fluctuation from the first phase stops at 200k updates, and we instead reward the tasks of the first phase (the basic task-set) as we did in the static treatment (all constant reward).

- **Harsh Changing Environment**: This treatment shows the effects of a continuing harsh changing environment on adaptation to the **expanded task set**. For the first 200k updates (stage 1), we alternate rewarding and **punishing** populations for performing the `EQU` task (Table 2). For the second stage of the experiment, starting at 200k updates, we add constant rewards for each of the new tasks of the **expanded task set**, at a 1.2-fold bonus to task execution. The environmental fluctuation from the first phase continues through the end of the experiment.
- **Harsh Quiescent Changing Environment**: In contrast to the harsh changing environment treatment, this treatment tests the abilities of populations initially evolved in a harsh changing environment to adapt to the **expanded task set**, but without active environmental fluctuation during the adaptation. For the first 200k updates (stage 1), we alternate rewarding and **punishing** populations, as in the Harsh Changing Environment above. For the second stage of the experiment, starting at 200k updates, we add constant rewards for each of the new tasks in the **expanded task set**, at a 1.2-fold bonus to task execution. The environmental fluctuation from the first stage stops at 200k updates, and we instead reward the basic tasks of the first phase as we did in the static treatment (all constant reward).

### Measuring Task Discovery and Task Performance

Task discovery and task performance are important measures not only of the adaptation of digital organisms to their local environment, but they also indicate the extent to which populations are more or less evolvable. Populations that are more evolvable should be able to acquire new tasks at a faster rate than less evolvable populations. If the evolvability of our populations is affected by evolution in a changing environment, then this effect should result in differential rates of task discovery and performance. Task discovery and performance together describe the exploration and exploitation of the environment by a population.

### Task Discovery

Task discovery represents the level of exploration of the fitness landscape. We measured task discovery by counting the number of unique non-ephemeral tasks that have been discovered by a population. Each task may be performed only once per organism, yielding a maximum task count of 3600 at any given time. We define a non-ephemeral task as one that is performed by at least than 0.1% of the population. Therefore, in order for a new task to be marked as discovered, it must be performed by at least 4 individuals at the time of sampling.

Once a task is discovered, it may not be un-discovered; task discovery counts will always increase monotonically. We measure **overall** task discovery by beginning to collect unique tasks starting at the beginning of the experiment. For the overall measurement, we count all possible tasks - the **expanded task-set** - even though not all tasks are rewarded in the first stage of the experiment. We also measure **post-reward** task discovery, where we begin counting new tasks from the beginning of the second stage of the experiment, once we have begun rewarding execution of the **expanded task set**. Task discovery can range anywhere from a minimum of zero tasks discovered, to a maximum of 77.

### Task Performance

In addition to counting the number of unique task discovered, we also measure task performance. We measure task performance by counting the total number of unique, non-ephemeral tasks that a population is performing at each sampling point. This measure represents the level of exploitation of the fitness landscape. This measure can range from 0 to a maximum of 77 task being performed by the population. This value will always be either equal to, or smaller than the number of tasks discovered, since a population can’t perform a task it hasn’t discovered yet.

### Experimental System

For all of the experiments described in this paper, we held the individual genomes at a fixed length of 121^2^ instructions, but tested the new genomes for mutations after each successful replication event at a substitution probability of 0.00075 per site.

We configured the Avida world to have local interactions on a toroidal grid that is 60-by-60 cells (3600 cells in total), and we seeded the initial populations with an ancestor that was previously evolved to perform `XOR` and `EQU` under a static reward. The genetic architecture for performing `XOR` and `EQU` is tightly intertwined in this ancestral organism, as it was evolved with no selective pressure for modularity.

## Statistical Methods

In experiments with digital organisms, it is fairly simple to perform so many replicates that questions of the meaning of significance arise. In order to paint a true statistical picture of the differences between our controls and experimental conditions, we limited our replicates to 50 for each experimental condition, and emphasize the effect size in all our statistical claims, in addition to reporting significance.

Most of the statistical techniques used in this paper are non-parametric, and focused on differentiating between sample distributions. In general, we applied Wilcoxon Rank-Sum tests [64] to distinguish between pairs of distributions, as well as Kruskal-Wallis [65] for identifying whether we could reject the null hypothesis of sameness between several different distributions. We assume all distributions are independent, and that compared distributions have similar shapes. In all situations where there were multiple comparisons of a given distributions, we applied Bonferonni corrections [66] before assessing statistical significance.

In certain cases, we report mean and median values of distributions. In these cases, we also report the standard deviation or 95% confidence intervals.

In specific cases, we also apply Spearman’s rank-order correlation coefficient *ρ* (or *r*_*s*_) [67] to measure correlations between data sets. In all cases, data points are matched from within a replicate.

All statistics were calculated using the SciPy [68] and Pandas [69] packages, in Python [70].

## Results and Discussion

Our experiments (detailed below) demonstrate that digital organisms that were evolved in changing environments differ substantially from those that evolved in static environments in a number of ways. These differences include the number of mutations that fix in the lineage from the ancestor (the “phylogenetic depth”), key metrics of their genetic architecture, and the presence of reservoirs of pseudogenes that change the nearby mutational landscape. These features represent adaptation to the larger regime of repeated environmental switching.

We also show that while harsh changing environments are better at promoting short-term adaptation to changing environments, benign environments produce populations that adapt more rapidly to entirely new sets of tasks. This result suggests that the selective pressures that promote short-term adaptation are not necessarily the same as those that promote long-term evolvability.

### Stage 1 - Cyclic Changing Environments

We begin by examining the characteristics of populations evolved in cyclic changing environments.

### Performance of EQU

Each population was seeded with organisms that performed both the `EQU` fluctuating task, and the `XOR` backbone task. We measured the execution of the `EQU` task, and observed that in the static control treatment, `EQU` is fixed in the population and remains so throughout the run. In contrast, we observed a periodic dip in the execution of `EQU` in the benign changing environment during the non-rewarded phase of the cycle, followed by a rapid recovery when rewards are reinstated. Finally, in the harsh treatment, we observed abrupt disappearance of `EQU` performance, followed by rapid recoveries, coinciding with phases of reward and punishment. As expected, these results suggest that the populations are responding to the selective pressures to perform `EQU` when it is rewarded, and to lose functionality when it is not rewarded or when it is punished.

### Evolutionary History and Population Structure

We then surveyed the evolutionary history and population structure of the evolving populations. Evolution in the harsh cyclic changing environment resulted in many more mutations fixing, and thus populations with substantially higher phylogenetic depth as compared to those evolved in static or benign environments. At each environmental shift, adaptive mutations rapidly swept and fixed in the populations. (Fig 3)

**Fig 3.**
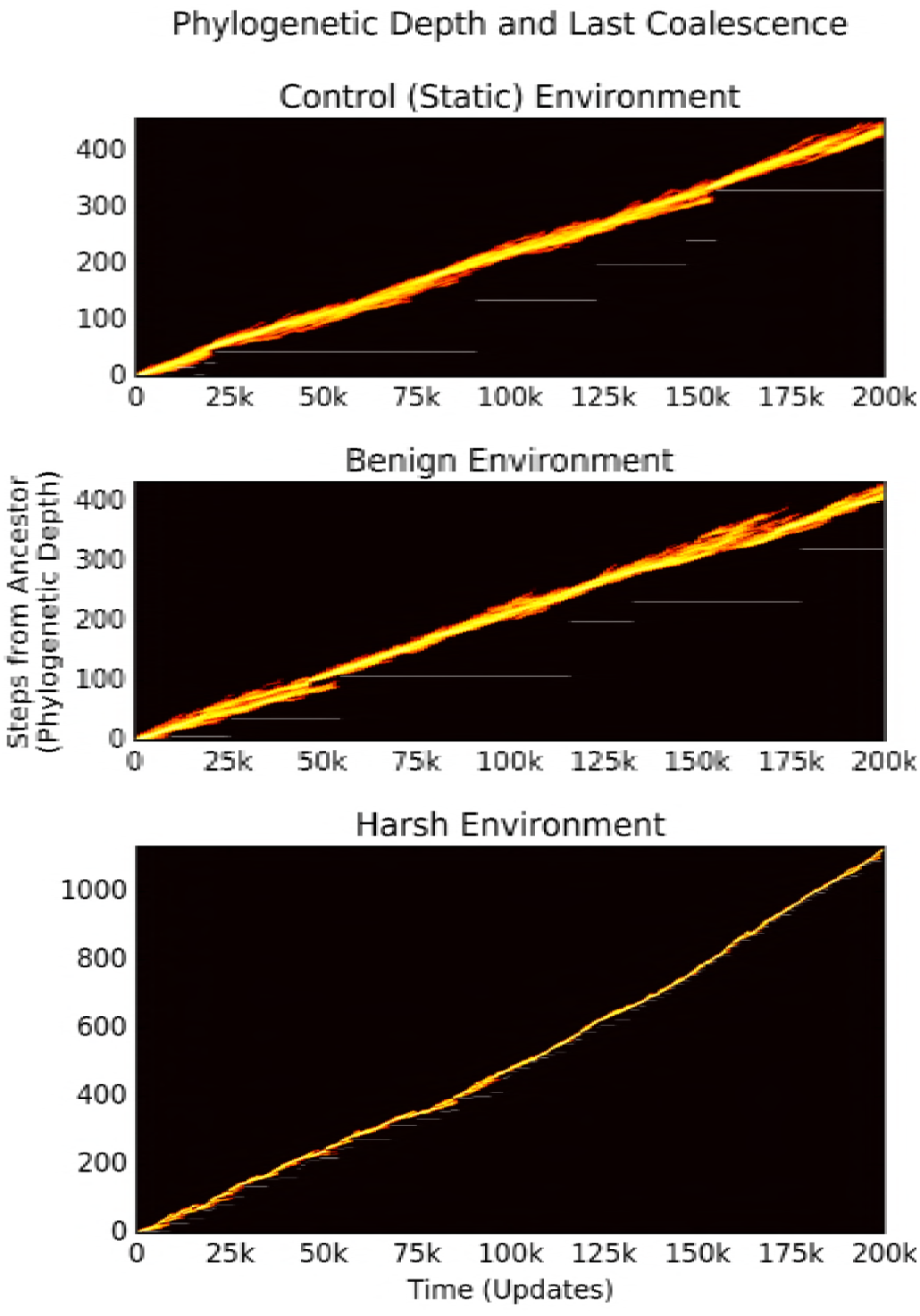
**Phylogenetic depth over time** of a sample population evolved in each of the three treatments of the cyclic changing environments. White horizontal lines mark the depth of the most recent common ancestor, and discontinuities in this line indicate that the most recent common ancestor has changed, and thus that a sweep occurred, or that a competing clade went extinct. The control treatments had a mean of 18 sweeps (STD=9.05), the benign treatments had a mean of 21 (STD=19.05), and the harsh treatments had a mean of 88 sweeps (STD=23.37). Note the difference in scales between y-axes: the control-evolved population has a maximum depth of 400 mutational steps from ancestor, while the harsh-evolved has upward of 1100.

The populations that evolved in the control and benign environments displayed more genetic diversity as compared to those evolved in the harsh cyclic environment, which underwent a bottleneck at each cycle shift (see Fig 5). Because a selective sweep reduces current diversity within a population, the smaller number of sweeps in the benign and control treatments led populations in them to have higher standing diversity for most of their evolutionary history than those populations from the harsh changing environment. Despite this higher standing diversity in the benign and control treatments, regions of low diversity are still evident in the genomes of these populations, implying purifying selection on the traits encoded at these sites (see Fig 4).

**Fig 4.**
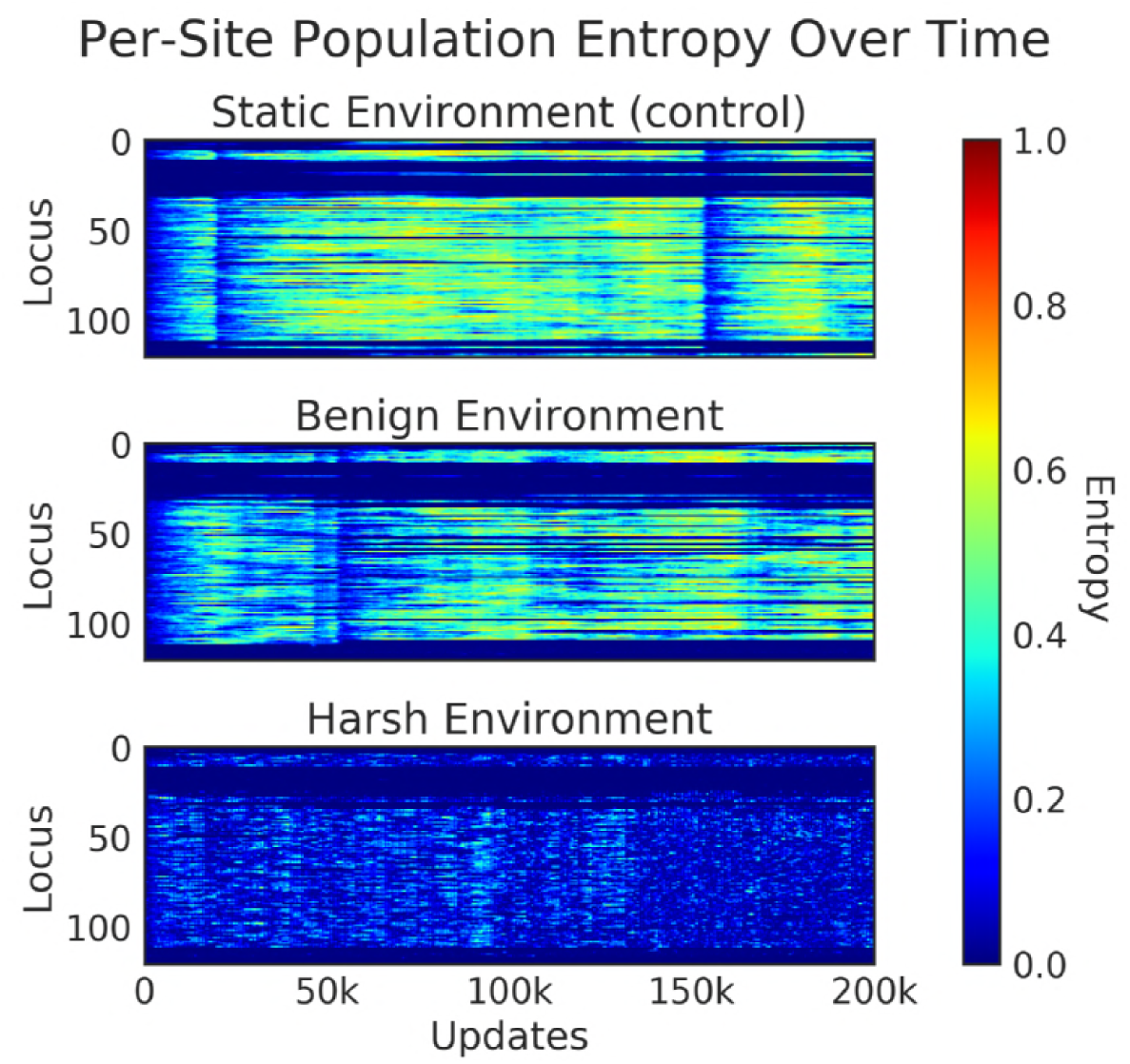
**Per-site entropy over time** of a representative sample population. Each vertical slice represents the per-site entropy of the population at each update by genetic locus. Hotter colors (red/orange/yellow) indicate greater diversity at this locus, while cooler colors (blues) indicate that a locus is more consistent across the population. These data indicates that in harsher changing environments, per-site diversity was much lower than in benign or static environments.

**Fig 5.**
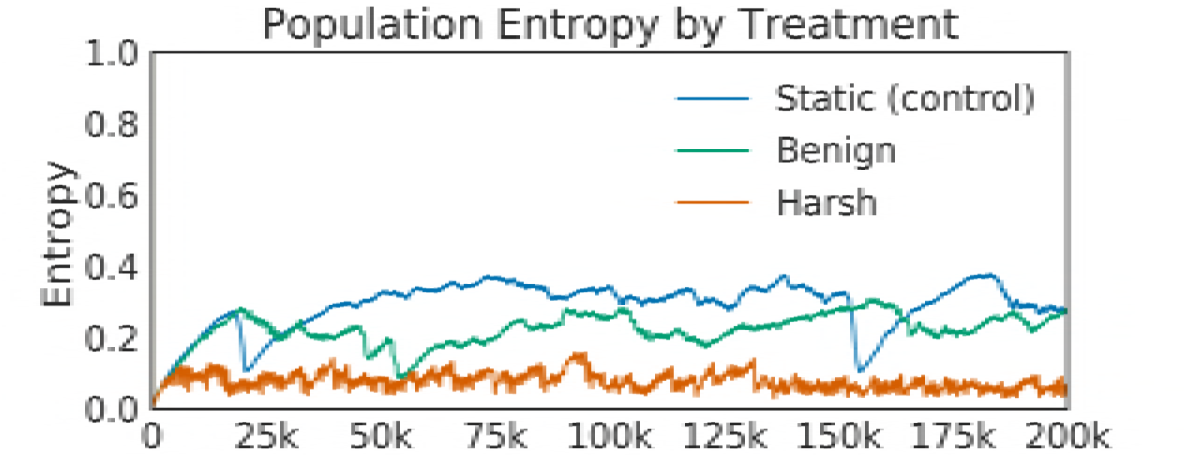
**Population entropy over time** of the representative sample population in Figure 4. Mean population entropy indicates the relative diversity of the population at any given time, while the per-site entropy (see Fig 4) shows where in the genomes the population diversity is located. These data indicate that in harsher changing environments, as in per-site diversity (above), overall population diversity was much lower than in benign or static environments.

### Genetic Architecture

The alternating selection in both benign and harsh changing environments results in qualitatively different architectural styles as compared with those genomes evolved in the static environment. The task arrangements evolved under both experimental treatments are much more scattered throughout the genome than in the control, which is tightly compacted. Specifically, the bulk of the sites responsible for performing the fluctuating task (EQU) did not overlap with the backbone task (`XOR`), except for a small core region, which represents portions of the tasks that are shared between `XOR` and `EQU`. That is, in the changing environment treatments, we see many more sites that only code for a single task, whereas in the static treatment, the majority of functional tasks sites code for both `XOR` and `EQU`. (See Figs 6, 7, and 8)

**Fig 6.**
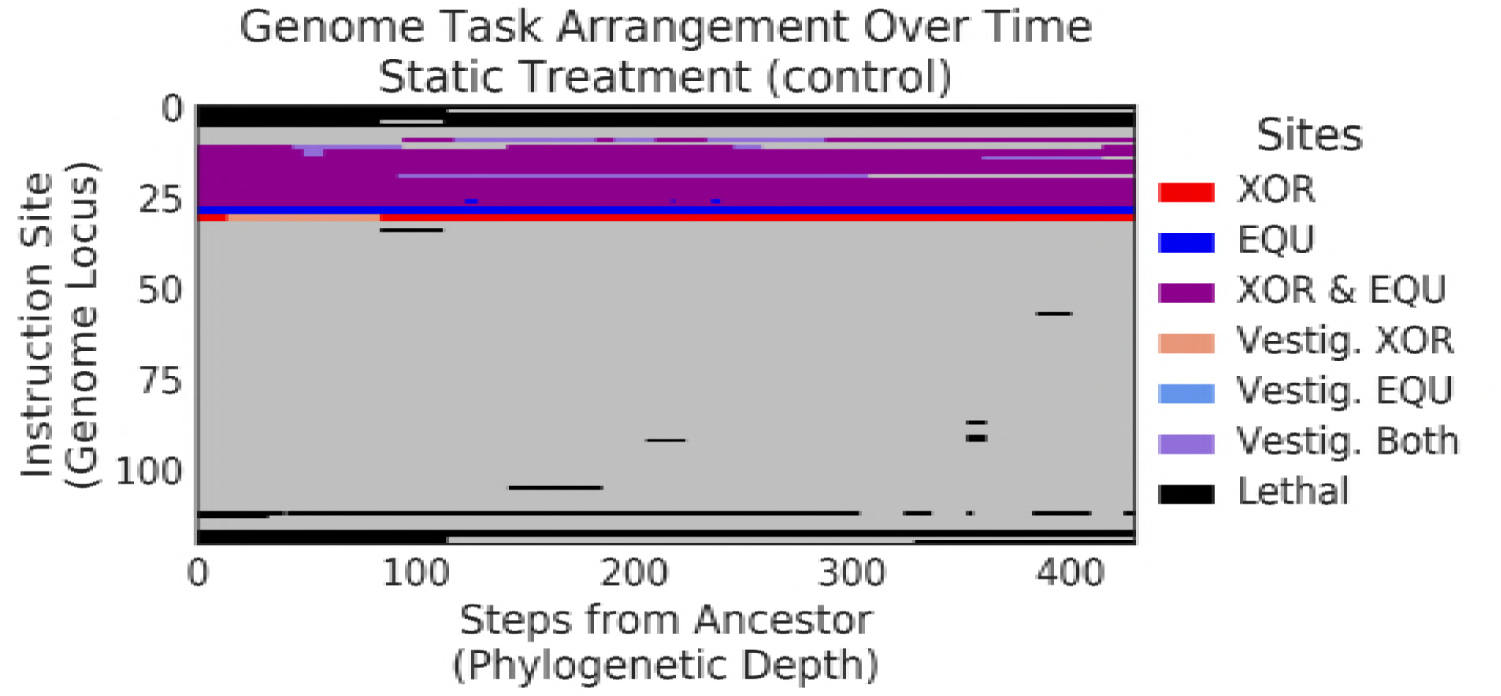
**Genetic architecture of `XOR` and `EQU` over time in static environment** for the final dominant genotype in a randomly selected replicate. Starting from the ancestor on the left, each vertical slice represents an organism along the line of descent to the final dominant. Positions along the Y-axis represent each genome locus; loci in an organism are colored based on the tasks that they currently (or previously) code for. Sites in **red** are active sites that code for the `XOR` task only, sites in **blue** code for the `EQU` task only, and **purple** sites code for both. Sites in black are critical for organism replication. Sites in the lighter colors (tan, light blue, lavender) represent vestigial sites, unchanged since they previously coded for `XOR` only, `EQU` only, or both tasks, respectively. As we proceed from left to right, we can see the evolutionary history of the final dominant genotype. `XOR` and `EQU` overlap almost completely throughout the run. This kind of genetic architecture is typical of purely directional selection, where a population has stabilized around a fitness peak.

**Fig 7.**
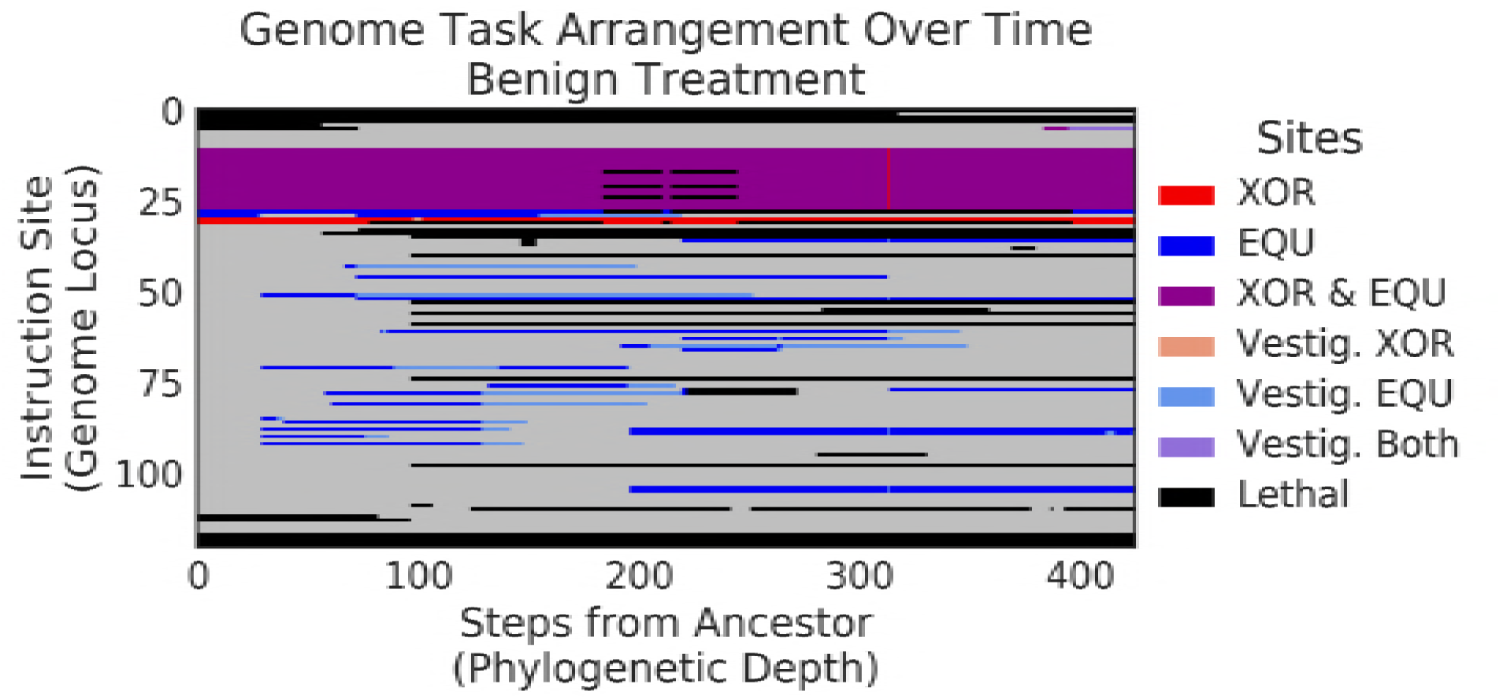
**Genetic architecture of `XOR` and `EQU` over time in benign environment** for the final dominant genotype in a randomly selected replicate. Proceeding from the left of each figure, each vertical slice represents an organism along the line-of-descent to the final dominant, and as in Figure 6, colors represent tasks performed by each genome locus. In this genome, `XOR` and `EQU` evolve to overlap much less than in the control. In contrast to the control (above), because fitness peaks are changing over time, more functional mutations are accepted.

**Fig 8.**
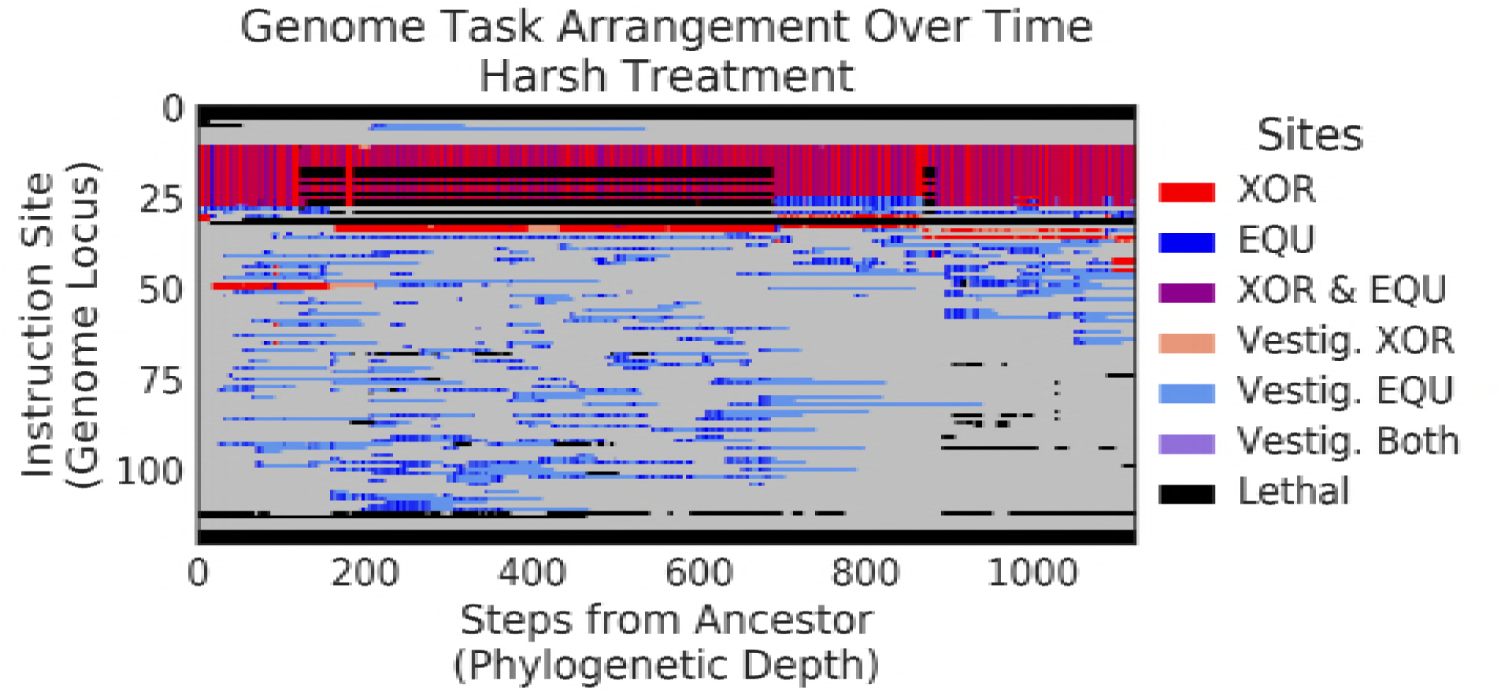
**Genetic architecture of `XOR` and `EQU` over time in harsh environment** for the final dominant genotype in a randomly selected replicate. Proceeding from the left of each figure, each vertical slice represents an organism along the line-of-descent to the final dominant, and as in Figures 6 and 7, colors represent tasks performed by each genome locus. In this genome, `XOR` and `EQU` evolve to overlap even less than in the control and benign treatments, with the `EQU`-only task sites becoming increasingly scattered throughout the genome. In contrast to both the control and benign environments (above), the harsh changes in selective pressure promote the adoption of many more mutations, and result in a much different genetic architecture.

In terms of site placement over time, functional task site locations in the control treatment did not change substantially over the course of the experiment. In the benign treatment, many more regions that performed the fluctuating task (`XOR`) were scattered throughout the genome, but site positions remained relatively fixed throughout the run after an initial adaptive phase. In the harsh treatment, however, not only were the active sites scattered, but the positions of active sites changed and proliferated wildly over time.

In addition to the variation in site placement, populations in the benign and harsh changing environment treatments had significantly more functional sites devoted to performing just the `EQU` task (Wilcoxon Rank Sum Test: Z = −5.57 and −6.96, respectively, p << 0.001). Interestingly, populations evolved in both the benign and harsh treatments also show development of a large reservoir of formerly functional, now vestigial, sites; that is, sites that remain unchanged from when they were previously active in performing a task, but were disabled by a mutation elsewhere and are thus now neutral. These vestigial pseudogene-like sites may be important for allowing the organisms to quickly re-adapt as the fluctuations in the environment restore the previously-rewarded functions. (Fig 9)

**Fig 9.**
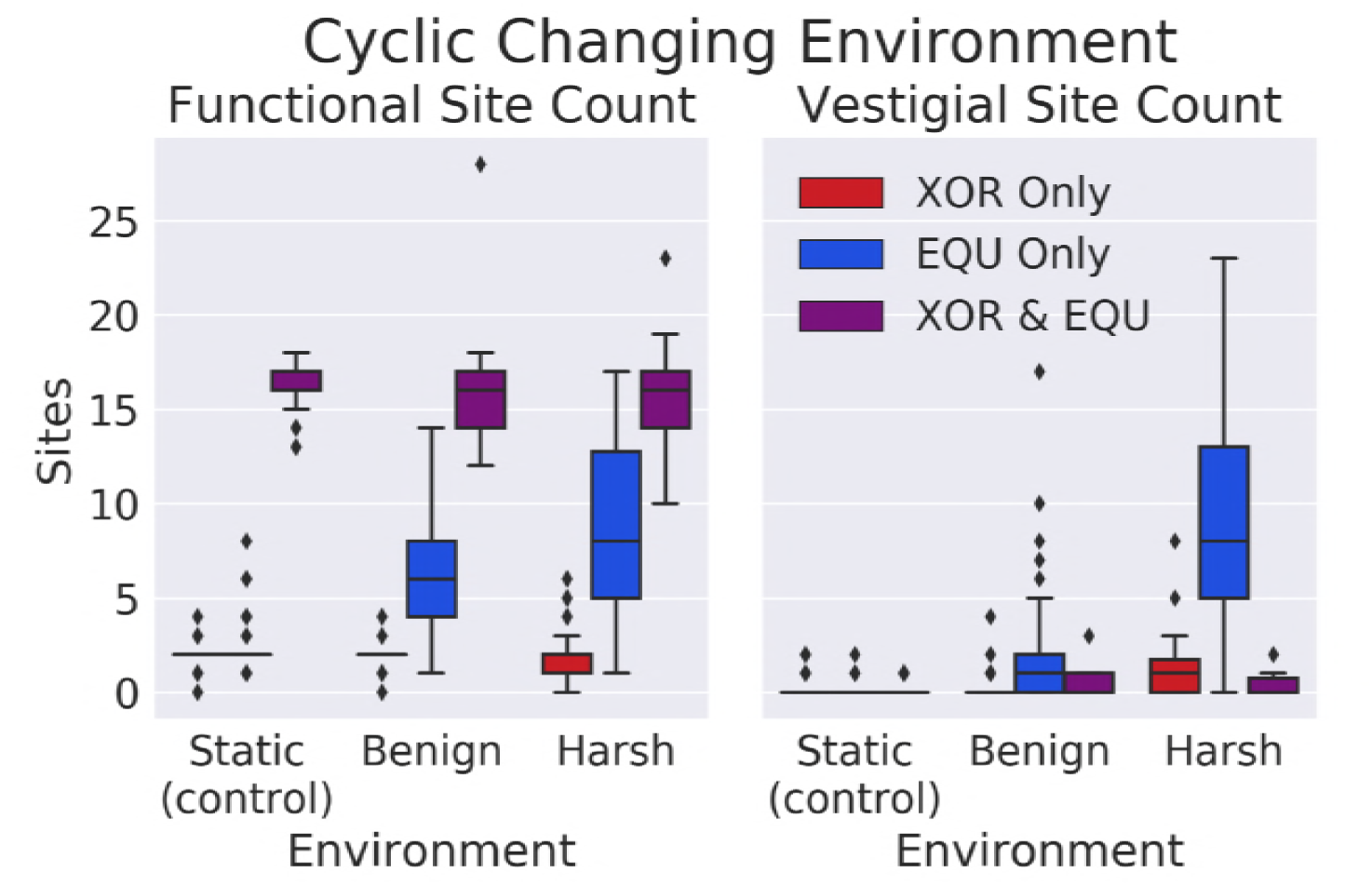
Number of functional and vestigial sites by treatment. Both the benign and harsh changing environments had significantly more sites devoted to performing only the `EQU` function (Wilcoxon Rank Sum Test: Z = −5.57 and −6.96, respectively, p << 0.001). The harsh environment has a significantly larger number of vestigial sites for the fluctuating (EQU) task compared to the benign treatment or control (Wilcoxon Rank-Sum Z = −6.57 and −8.33, p << 0.001).

### Nearby Mutational Landscape

In order to identify the role that these longer task footprints and pseudogene-like structures play, we performed a survey of the single-step mutational neighborhood surrounding the most abundant genotype present at the end of the experiment for each replicate population. Each neighborhood contained 3,025 distinct mutants (121 loci with 25 possible mutations per locus) in each of the 50 replicates per treatment, for a total of nearly 450,000 mutants surveyed. We measured the fraction of mutants that lost each of the rewarded tasks. (Fig 10).

**Fig 10.**
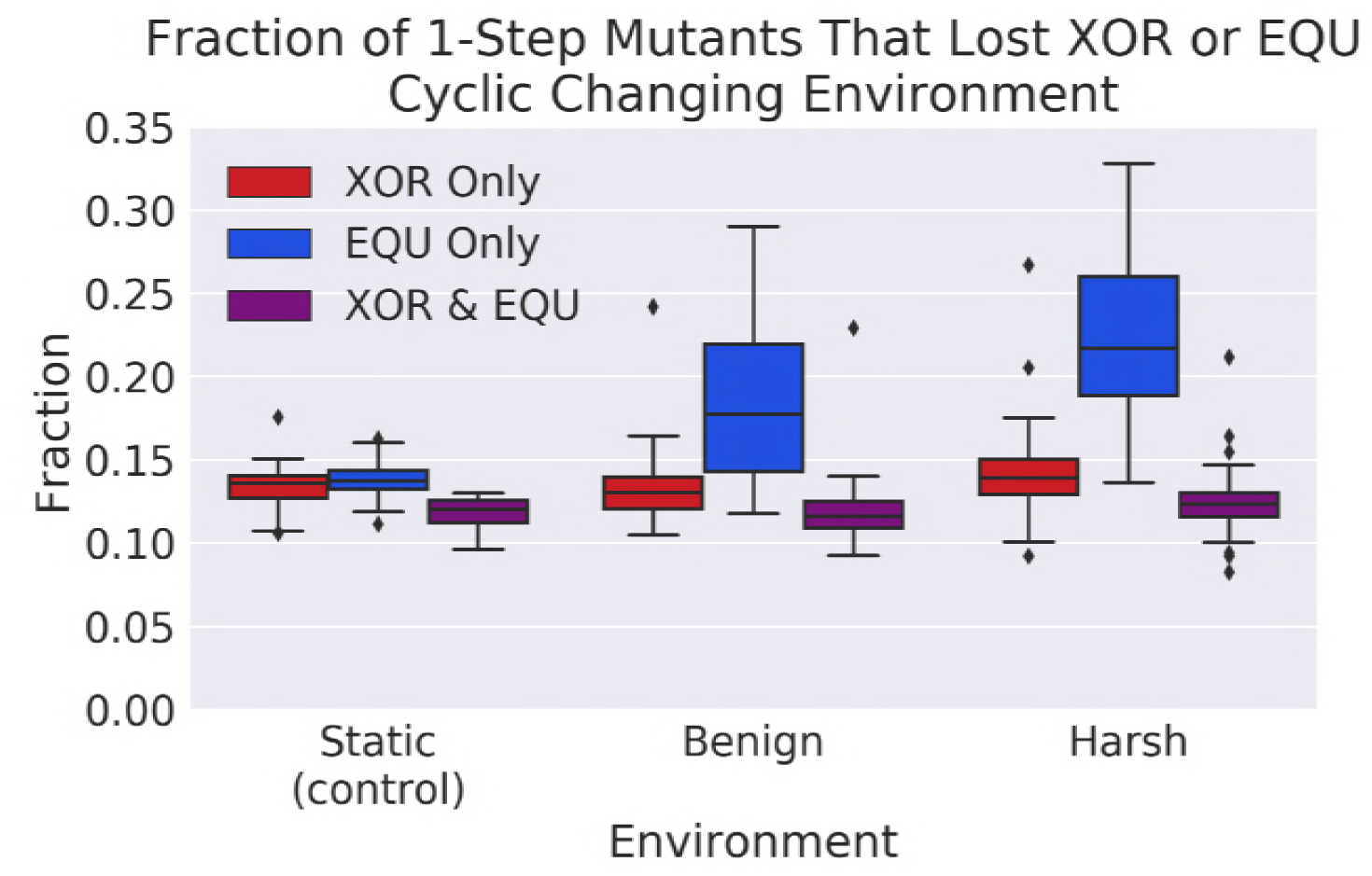
**A survey of the single-step mutational neighborhood** around organisms that performed the fluctuating task. Note that in both the benign and harsh treatments, there were significantly more mutants that lost the `EQU` task as compared to the control (Wilcoxon Rank Sum Test: Z = −5.46 and −7.80 respectively, p << 0.001). This result indicates that it was easier for the organisms in both treatments to turn off the `EQU` task in response to one mutation.

We found that in both the benign and harsh treatments, there were many more mutations that resulted in loss of the fluctuating task as compared to the control (Wilcoxon Rank Sum Test: Z = −5.46 and −7.80 respectively, p << 0.001). An increase in task loss in the harsh treatment is to be expected, but why would the benign treatment lose `EQU` nearly as easily as the harsh treatment? One possibility is selective pressure to lose the task. There is no explicit pressure for task loss, merely an absence of reward. Even so, there is certainly an implicit penalty for performing a complex task for which there is no reward.

Another possibility is drift. Indeed, in Figure 2, we observe a steady downward trend in execution of `EQU` when rewards are withdrawn. Then, as the reward returns, new mutations are applied that reactivate the task, and overall performance recovers quickly. This pattern of loss and regain would, over time tend, to increase the length of the task. Indeed, as noted in Figure 8, there is a rapid increase in task length as `EQU` is cyclically lost and regained.

**Fig 2.**
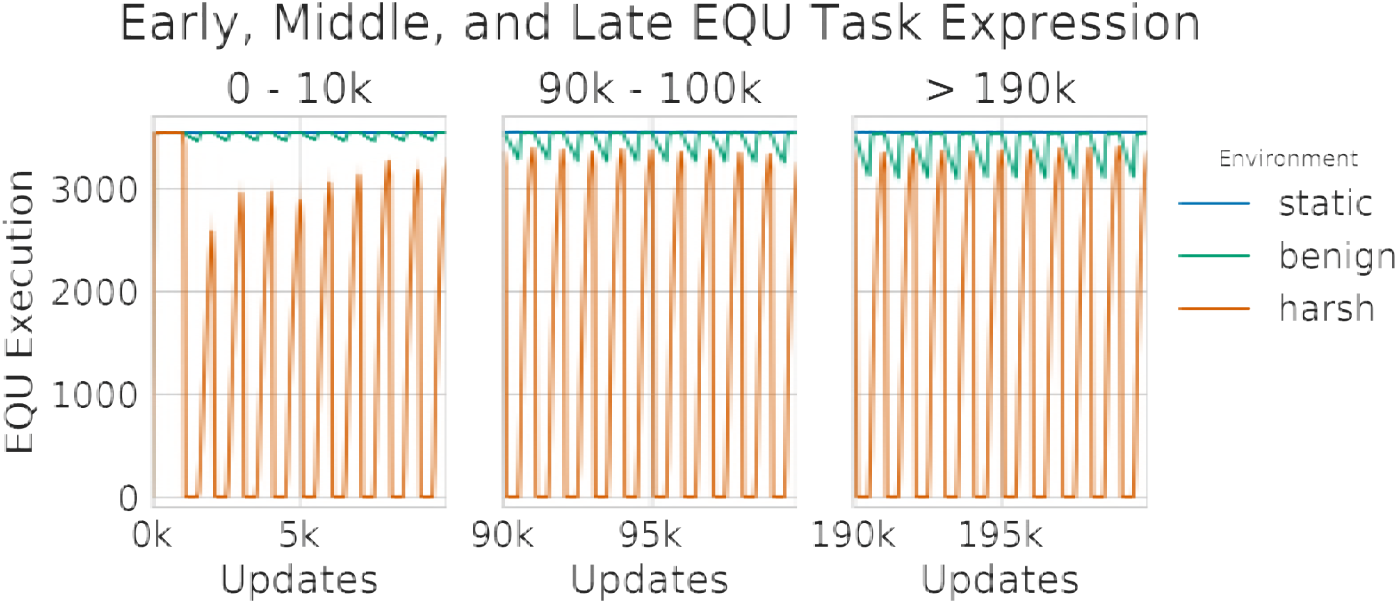
Number of organisms performing EQU task. We measured the execution of the `EQU` task in all treatments. In the benign treatment, we observed increasing periodic dips in execution that coincide with phases of non-reward as the experiments progressed. In the harsh treatment, we observed adaptation, resulting in abrupt disappearance of `EQU` in the punishment phase, followed by rapid recovery of `EQU` performance during the reward phase. This result suggests that, over time, populations became more apt at rapidly gaining and losing `EQU` as a response to changing selective pressures.

However, is increased task length enough to account for increased task vulnerability to mutation? In order to begin to address this question, we constructed a linear model relating the task length of each task with the fraction of mutants that lost each of the tasks. We discovered a strong relation between the number of functional sites and the number of task-losing mutants for the `EQU` task, both alone, and overlapping with `XOR`, such that we could predict approximately 77% and 63% of the variation in task loss, respectively (Fig 11, see Tables 3 and 4). We also found a weaker, but still significant relationship between the number of `XOR`-only functional sites and loss of the `XOR` task, such that we could predict approximately 22% (See Table 5). This result confirms our intuition that the longer the task, the more targets there are for mutation to disable the task.

**Table 3.**
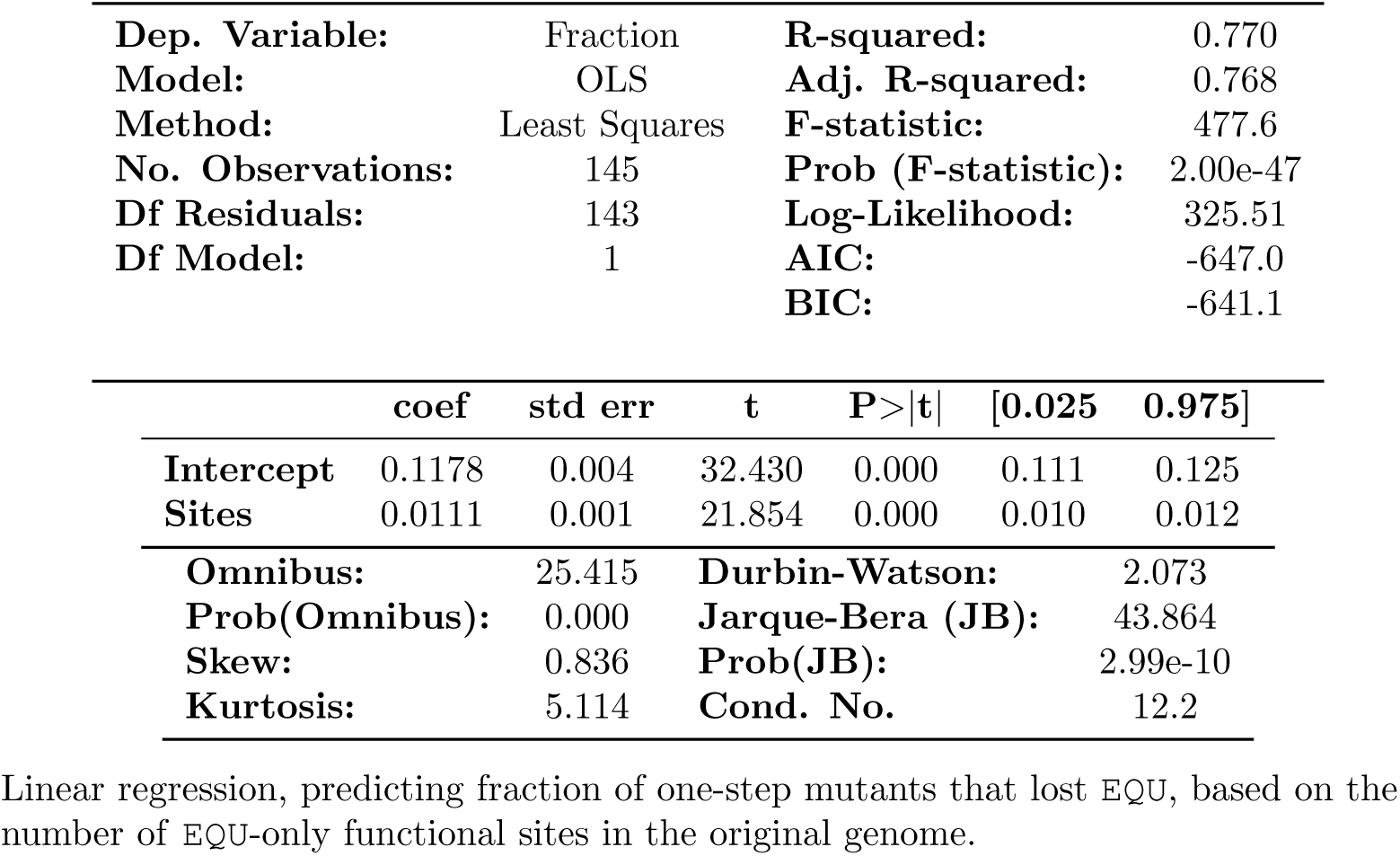
OLS Regression Results - Fraction of mutants losing EQU vs number of EQU-only functional sites.

|  |  |  |  |
| --- | --- | --- | --- |
| Dep. Variable: | Fraction | R-squared: | 0.770 |
| Model: | OLS | Adj. R-squared: | 0.768 |
| Method: | Least Squares | F-statistic: | 477.6 |
| No. Observations: | 145 | Prob (F-statistic): | 2.00e-47 |
| Df Residuals: | 143 | Log-Likelihood: | 325.51 |
| Df Model: | 1 | AIC: | -647.0 |
|  |  | BIC: | -641.1 |

|  | coef | std err | t | P>\|t\| | [0.025 | 0.975] |
| --- | --- | --- | --- | --- | --- | --- |
| Intercept | 0.1178 | 0.004 | 32.430 | 0.000 | 0.111 | 0.125 |
| Sites | 0.0111 | 0.001 | 21.854 | 0.000 | 0.010 | 0.012 |
| Omnibus: | 25.415 | Durbin-Watson: | 2.073 |
| Prob(Omnibus): | 0.000 | Jarque-Bera (JB): | 43.864 |
| Skew: | 0.836 | Prob(JB): | 2.99e-10 |
| Kurtosis: | 5.114 | Cond. No. | 12.2 |

Linear regression, predicting fraction of one-step mutants that lost EQU, based on the number of EQU-only functional sites in the original genome.

**Table 4.**
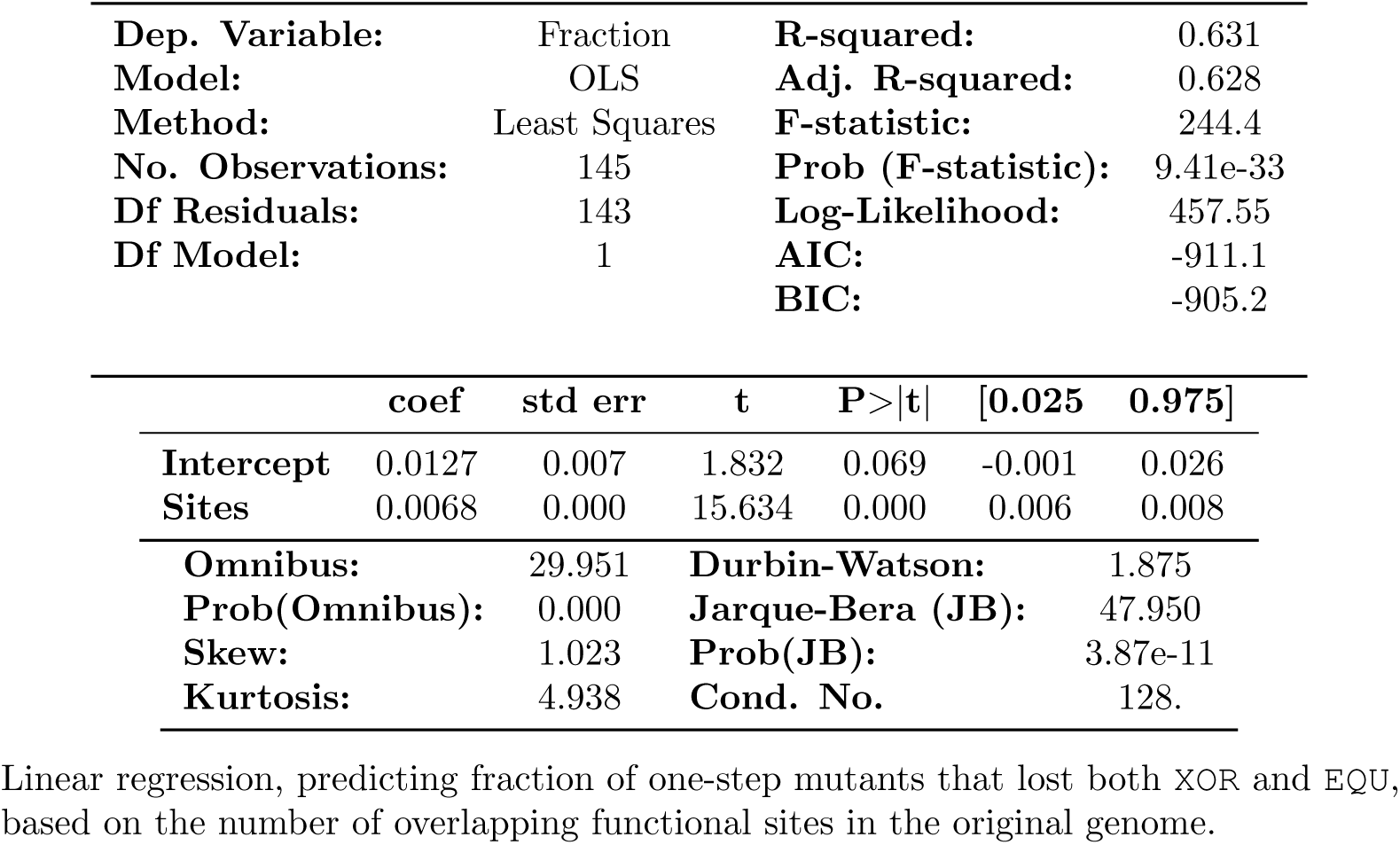
OLS Regression Results - Fraction of mutants losing EQU and XOR vs number of overlapping functional sites.

|  |  |  |  |
| --- | --- | --- | --- |
| Dep. Variable: | Fraction | R-squared: | 0.631 |
| Model: | OLS | Adj. R-squared: | 0.628 |
| Method: | Least Squares | F-statistic: | 244.4 |
| No. Observations: | 145 | Prob (F-statistic): | 9.41e-33 |
| Df Residuals: | 143 | Log-Likelihood: | 457.55 |
| Df Model: | 1 | AIC: | -911.1 |
|  |  | BIC: | -905.2 |

|  | coef | std err | t | P>\|t\| | [0.025 | 0.975] |
| --- | --- | --- | --- | --- | --- | --- |
| Intercept | 0.0127 | 0.007 | 1.832 | 0.069 | -0.001 | 0.026 |
| Sites | 0.0068 | 0.000 | 15.634 | 0.000 | 0.006 | 0.008 |
| Omnibus: | 29.951 | Durbin-Watson: | 1.875 |
| Prob(Omnibus): | 0.000 | Jarque-Bera (JB): | 47.950 |
| Skew: | 1.023 | Prob(JB): | 3.87e-11 |
| Kurtosis: | 4.938 | Cond. No. | 128. |

Linear regression, predicting fraction of one-step mutants that lost both XOR and EQU, based on the number of overlapping functional sites in the original genome.

**Table 5.**
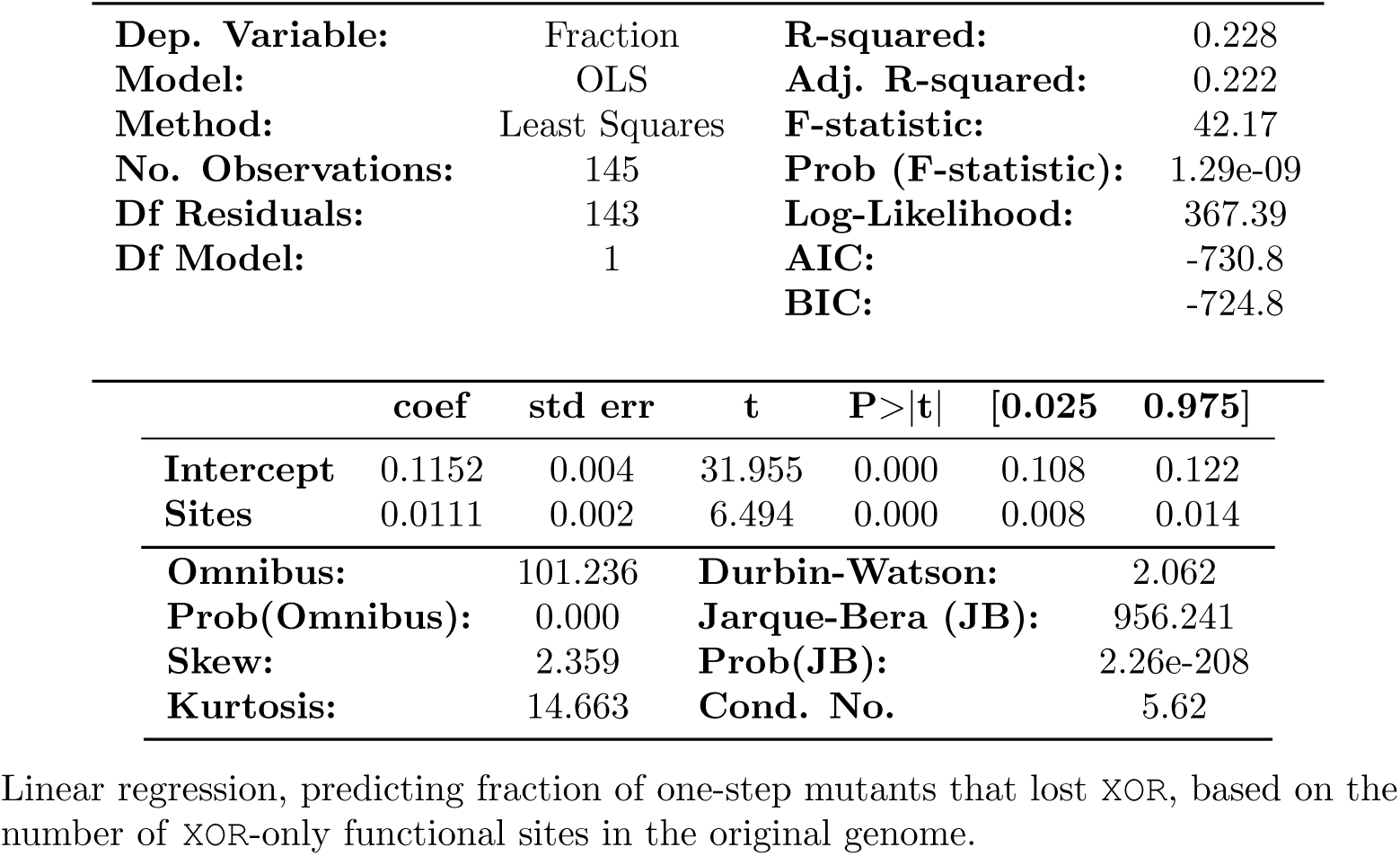
OLS Regression Results - Fraction of mutants losing XOR vs number of XOR-only functional sites.

|  |  |  |  |
| --- | --- | --- | --- |
| **Dep. Variable:** | Fraction | **R-squared:** | 0.228 |
| **Model:** | OLS | **Adj. R-squared:** | 0.222 |
| **Method:** | Least Squares | **F-statistic:** | 42.17 |
| **No. Observations:** | 145 | **Prob (F-statistic):** | 1.29e-09 |
| **Df Residuals:** | 143 | **Log-Likelihood:** | 367.39 |
| **Df Model:** | 1 | **AIC:** | -730.8 |
|  |  | **BIC:** | -724.8 |

|  | coef | std err | t | P>\|t\| | [0.025 | 0.975] |
| --- | --- | --- | --- | --- | --- | --- |
| **Intercept** | 0.1152 | 0.004 | 31.955 | 0.000 | 0.108 | 0.122 |
| **Sites** | 0.0111 | 0.002 | 6.494 | 0.000 | 0.008 | 0.014 |
| **Omnibus:** | 101.236 | **Durbin-Watson:** | 2.062 |
| **Prob(Omnibus):** | 0.000 | **Jarque-Bera (JB):** | 956.241 |
| **Skew:** | 2.359 | **Prob(JB):** | 2.26e-208 |
| **Kurtosis:** | 14.663 | **Cond. No.** | 5.62 |

Linear regression, predicting fraction of one-step mutants that lost XOR, based on the number of XOR-only functional sites in the original genome.

**Fig 11.**
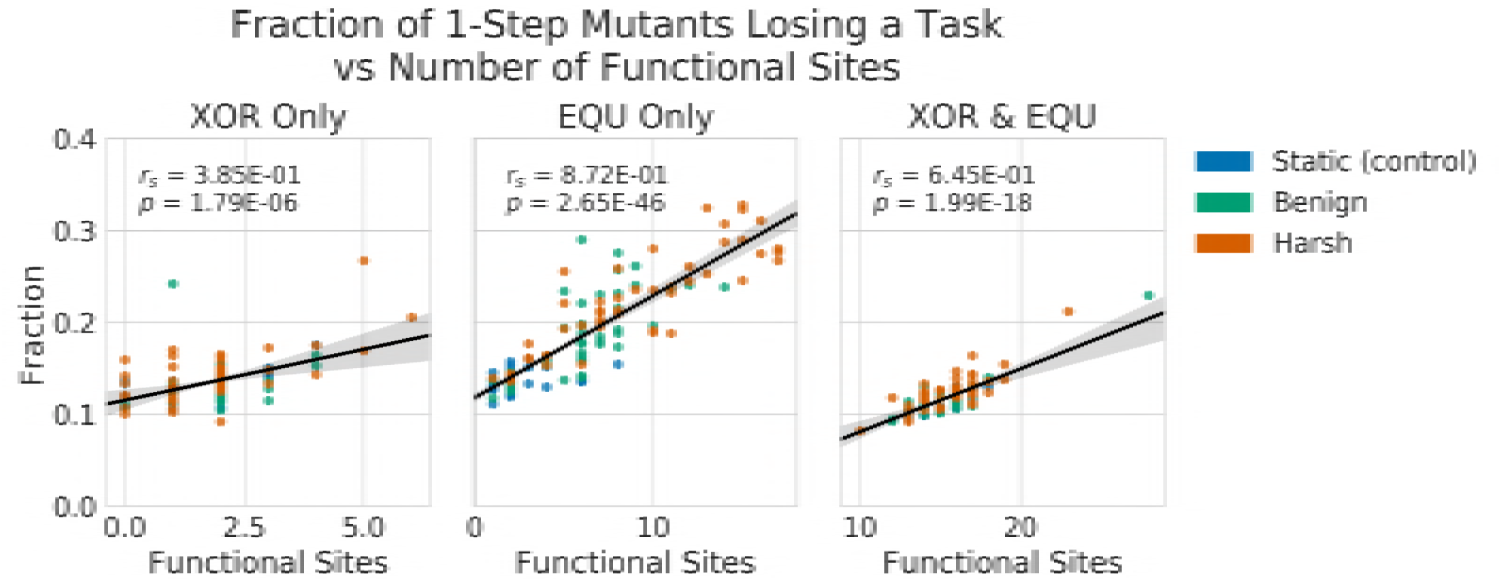
**Correlation between task length and mutational task loss** in the 1-step neighborhood across all treatments. Note the strong correlation between the length of the `EQU` task and the fraction of mutants that lost `EQU` (Spearman’s Rho: *r*_*s*_= 8.72, p << 0.001). Further, consider the weaker correlation between `XOR` task length, and fraction of mutants that lost `XOR`. These data suggest that `EQU` is even less robust to mutation compared to `XOR` than can be accounted for by task length alone.

Further, the lower correlation between length and task loss for the `XOR` suggests that it is not only task length, but some other architectural feature that makes the `XOR` task more robust to mutation, and the `EQU` task more fragile. Even so, the question of what kinds of architectural features account for this differential robustness remains open.

We then measured the proportion of second step mutants that regained `EQU` after having lost it in the single-step survey. We found that changing environments shifted the populations’ position in the mutational landscape, such that when a task that was lost due to mutation, that task could be regained via one or two additional mutations elsewhere. That is, once a mutation caused the loss of a task, a different mutation could reactivate the task. (Fig 12).

**Fig 12.**
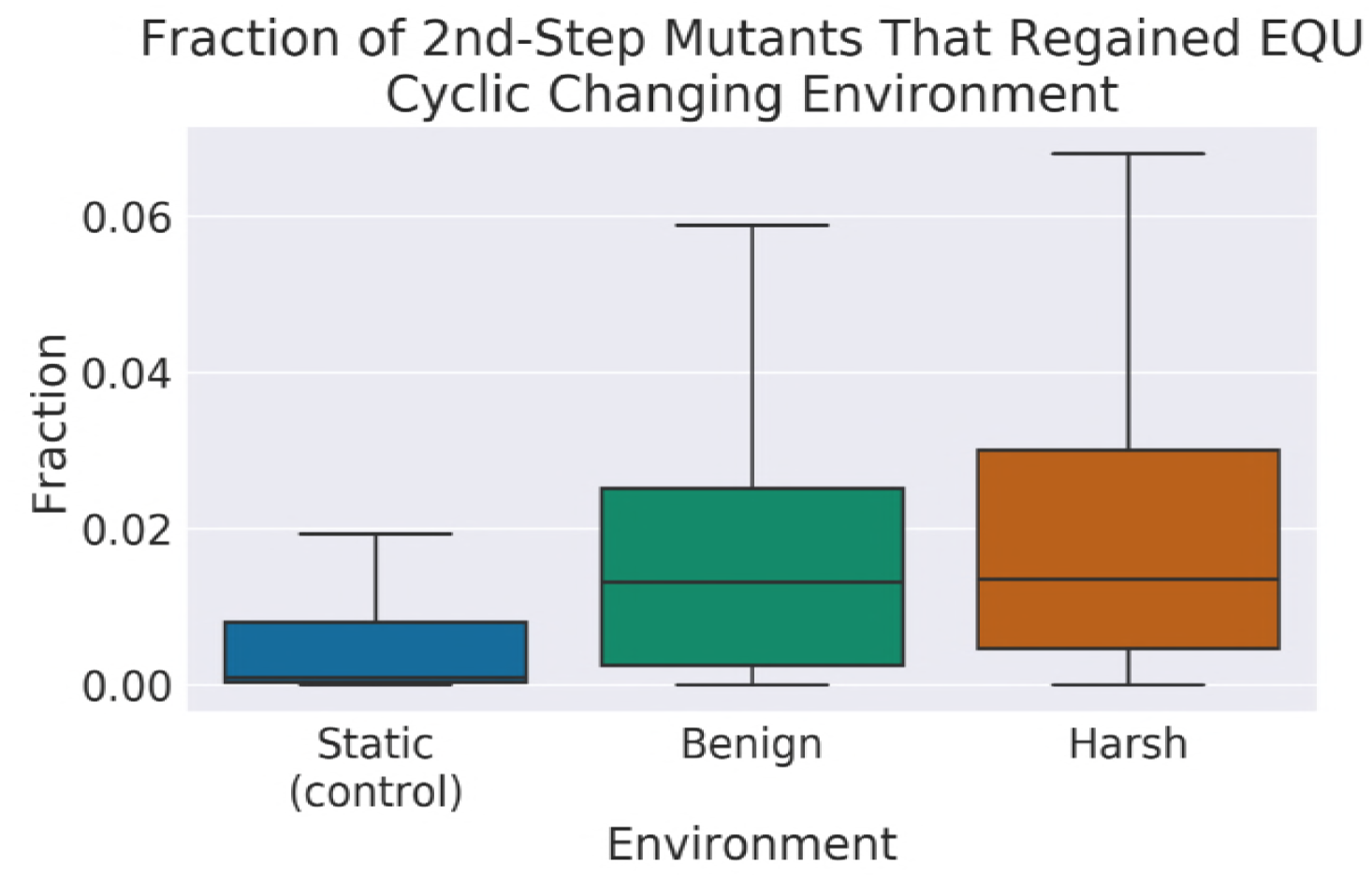
**A survey of the two-step mutational neighborhood** of the organisms that lost `EQU` function in the one-step survey. We found that in both the harsh and benign treatments, there were significantly more organisms that regained function in response to mutation than the control. (Wilcoxon Rank Sum Test: Z = −47.9 and −57.82 respectively, p << 0.001). This result indicates that it was easier for the organisms in both fluctuating environments to regain the task in response to one additional, non-reversion mutation.

We speculate that this effect is due to the availability of reservoirs of formerly vestigial sites. How such reservoirs might perform these functions remains an open question. New mutations may either re-enable the old functional sites, or recruit vestigial functionality to perform the task elsewhere. Potentially, these vestigial sites are not altogether dormant at all. They might individually appear vestigial in the context of a single knockout survey, but they might also be related to other sites in a network of backup functionality that becomes activated in response to mutation. More research is needed to explore what role these feature play.

As an overall measure of neutral exploration, we also measured the proportion of non-deleterious mutants in the nearby fitness landscape - the Genomic Diffusion Rate. We found that this proportion remained approximately the same between all treatments (Kruskal-Wallis: H(2) = 1.44, p = 0.49). However, we found that the Phenotypic Diffusion Rate, the proportion of these mutants with different (and potentially adaptive) phenotypes, increased in the changing environment treatments as compared to the controls (Wilcoxon Rank Sum Test: Z = −8.02, −8.39, respectively, p << 0.001). In this way, the organisms from the changing environment treatments have an advantage over organisms from the control runs in the short-term evolvability of the fluctuating task. This result is consistent with real adaptation, not only to resources in their local environment, but a direct adaptation to the environmental change. (Fig 13)

**Fig 13.**
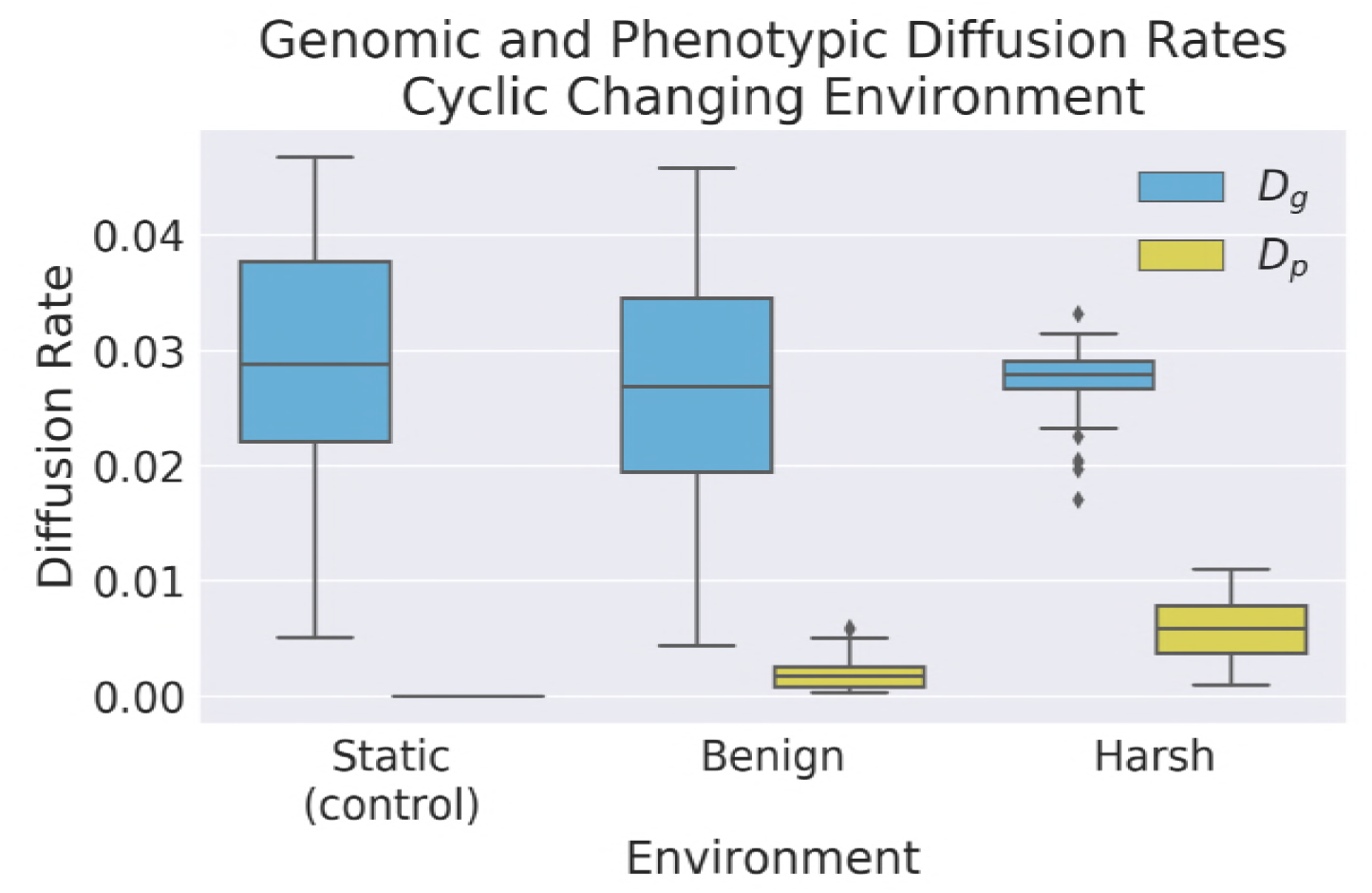
**Genomic and phenotypic diffusion rates**, showing the probabilities of producing offspring that are genotypically (*D*_*g*_) or phenotypically (*D*_*p*_) distinct from the parent, while not reducing fitness. Note that while overall neutral exploration capacity remains relatively stable between treatments (Kruskal-Wallis: H(2) = 1.44, p = 0.49), phenotypic exploration capacity is increased in both treatments, but especially in the Harsh treatment. (Wilcoxon Rank Sum Test: Z = −8.02, −8.39, respectively, p << 0.001). This result is consistent with changing environments promoting the phenotypic evolvability of populations.

What might account for this adaptation? Similar to the relationship between the number of functional sites of a task, and the number of single-step mutants that lost that task (see Fig 11), we hypothesize that the reacquisition of tasks in the 2nd-step survey may be mediated by the amount of useful task material present in the genome. We performed a multiple linear regression, predicting the mean fraction of mutants that regained `EQU`, by the number of functional and vestigial sites contained in the original genome (see Table 6 and Fig 14).

**Table 6.**
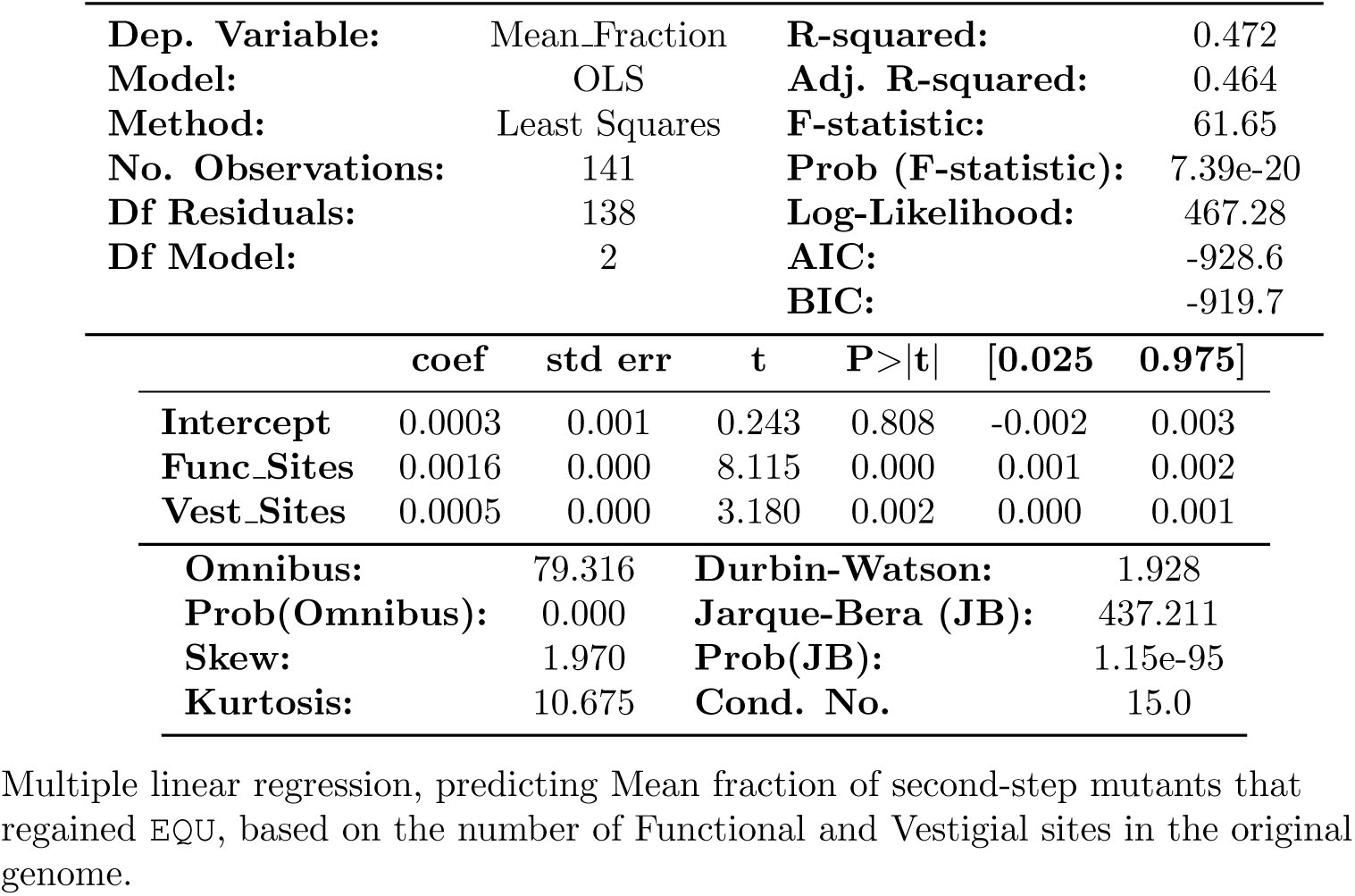
OLS Regression Results - Mean Fraction of Mutants Regained EQU vs Number of Functional and Vestigial Sites.

|  |  |  |  |
| --- | --- | --- | --- |
| **Dep. Variable:** | Mean_Fraction | **R-squared:** | 0.472 |
| **Model:** | OLS | **Adj. R-squared:** | 0.464 |
| **Method:** | Least Squares | **F-statistic:** | 61.65 |
| **No. Observations:** | 141 | **Prob (F-statistic):** | 7.39e-20 |
| **Df Residuals:** | 138 | **Log-Likelihood:** | 467.28 |
| **Df Model:** | 2 | **AIC:** | -928.6 |
|  |  | **BIC:** | -919.7 |

|  | coef | std err | t | P>\|t\| | [0.025 | 0.975] |
| --- | --- | --- | --- | --- | --- | --- |
| **Intercept** | 0.0003 | 0.001 | 0.243 | 0.808 | -0.002 | 0.003 |
| **Func_Sites** | 0.0016 | 0.000 | 8.115 | 0.000 | 0.001 | 0.002 |
| **Vest_Sites** | 0.0005 | 0.000 | 3.180 | 0.002 | 0.000 | 0.001 |
| **Omnibus:** | 79.316 | **Durbin-Watson:** | 1.928 |
| **Prob(Omnibus):** | 0.000 | **Jarque-Bera (JB):** | 437.211 |
| **Skew:** | 1.970 | **Prob(JB):** | 1.15e-95 |
| **Kurtosis:** | 10.675 | **Cond. No.** | 15.0 |

Multiple linear regression, predicting Mean fraction of second-step mutants that regained `EQU`, based on the number of Functional and Vestigial sites in the original genome.

**Fig 14.**
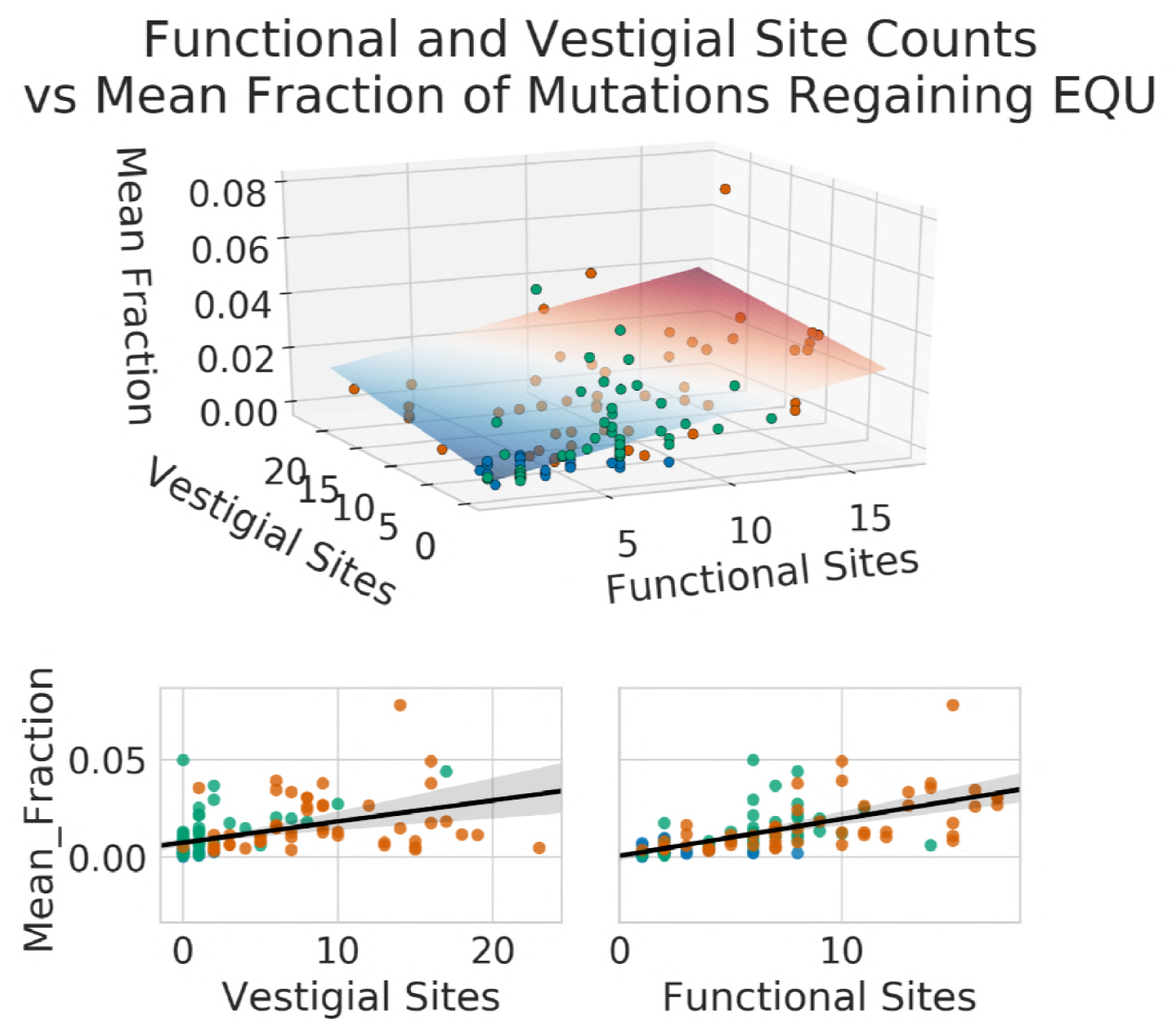
**Multiple linear regression predicting mean fraction of second-step mutants** from the number of functional and vestigial sites in the original organism. See Table 6.

We can predict approximately 47% of the variation in mean number of second step mutants that regained `EQU` based on the number of functional and vestigial sites. Most of the variation is predicted by the number of functional sites, though vestigial sites do contribute a small amount. This result is consistent with the role of task length, and thus the number of informational sites, being important for regaining task function. We could not, however, account for all variation, indicating that there are other factors, possibly in robustness or modular architecture of tasks, that are important to this kind evolvability.

### Stage 2 - Long-Term Evolvability

#### Task Discovery

Task discovery is an important measure of long-term evolvability in that it quantifies the ability of populations to explore and adapt to entirely new environments. We measured task discovery in each of the changing environment treatments.

#### Benign changing environments outperform harsh environments in task discovery

We found that once we began rewarding the **expanded task set**, populations evolving in harsh changing environments discovered many fewer tasks that those evolving in benign changing environments (Wilcoxon Rank Sum Test: Z = 2.75, p < 0.01) (Fig 15). We hypothesize that this effect is due to the relative differences in the strength of selection between the harsh changing environment and the directional selection toward the **expanded task set**.

**Fig 15.**
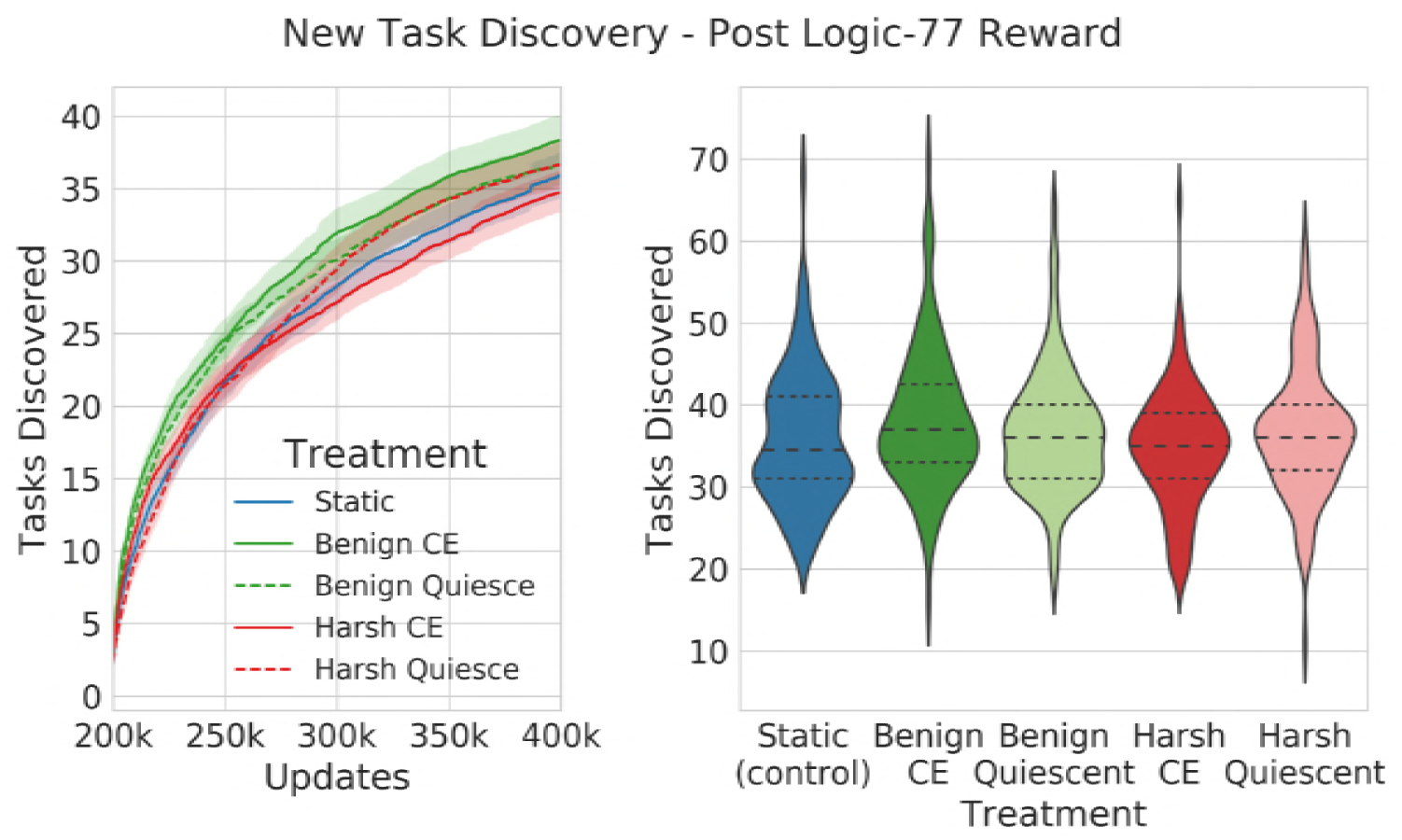
**Number of new expanded task set tasks discovered post-reward**. The left plot shows a time-series of the number of non-ephemeral tasks discovered by populations by each treatment. The right plot shows the number of tasks discovered at the end of the experiments. While the individual values overlap their neigbors, the top and bottom-most treatments (Benign and Harsh) are significantly different from each other (Wilcoxon Rank Sum Test: Z = 2.75, p < 0.01).

In the harsh changing environments, the selective pressure to gain or lose the fluctuating tasks represents up to a 2^6^-fold swing (between a × 2^5^ penalty and a × 2^5^ bonus) over the course of a single cycle, whereas the **expanded task set** can individually only offer a 1.2-fold bonus to execution speed. Thus, those organisms that promptly gain or lose a fluctuating task are more likely to survive, regardless of whether or not they have gained one of the new **expanded task-set** tasks. Thus pressures to gain and lose the fluctuating tasks are much stronger than the pressure to acquire new **expanded task-set** tasks, thereby depressing the rate at which they are found.

In contrast, the benign environment experiences a weaker strength of selection for `EQU` task gain and loss, in the form of a maximum 2^5^-fold bonus directional selection pressure to gain the tasks, and no direct pressure to lose the task. Thus, when compared to the harsh treatments, the fraction of the total selective pressure for gaining new expanded tasks is greater in the benign treatment. This increased pressure, plus the benefit of an increased exploration rate conveyed by the benign changing environment, result in a higher overall task discovery rates.

Interestingly, in the harsh quiescent treatment (HarshQuiescent) beginning in stage 2, we saw that task exploration recovered and achieved a comparable level to the control (Wilcoxon Rank Sum Test: Z = −0.91, p = 0.37). What could account for this recovery?

One possibility is that the introduction of the new tasks provided sufficient selective pressure to cause the increase in the discovery rate. In this case, we would expect to see a similar increase in task exploration in the harsh changing environment treatment (HarshCE).

Another possibility is that the alternating environment in the first part of the experiment created a diversity disadvantage in populations in those treatments. If this were the case, we would expect HarshQuiescent’s task discovery to initially grow more slowly than the control, which would have suffered from no such disadvantage. Then, as diversity recovered, we would expect to see task discovery grow at comparable rates.

Finally, there is the possibility is that the alternating selection regime was directly responsible for depressing task exploration. In this case, once we stopped alternating task rewards, we would expect to see a significant difference in task discovery rates between the HarshQuiescent and HarshCE treatments.

Indeed, we found that the HarshQuiescent treatment has a much higher task discovery rate than the HarshCE treatment. This result is inconsistent with the hypothesis that the task rewards alone account for the recovery of the HarshQuiescent. Instead, this result is consistent with the possibility of a direct negative effect from the continuing alternating selection. We also found that there was a lag in task discovery compared to the control. These data suggest that there was, at least initially, some population-level disadvantage occurring in the HarshQuiescent populations. We also observed that after the initially slow recovery phase, the quiescent treatment rapidly increased its task discovery rate, and exceeded that of the control. This outcome is consistent with a recovery of diversity, plus, potentially some lingering architectural advantage for finding new tasks.

### Harsh changing environments drive populations across the mutational landscape

In the first part of the experiments, despite the **expanded task set** not being rewarded, both changing-environment treatments (BenignCE and HarshCE) discovered more new tasks than the control (Wilcoxon Rank Sum Test: Z = −5.75 and −11.15 respectively, p << 0.001). The harsh treatment in particular discovered substantially and significantly more **expanded task set** tasks than either the benign treatment (Wilcoxon Rank Sum Test: Z = −8.0, p << 0.001) or the control, despite these tasks not being rewarded (Fig 16). We speculate that this effect may be due to the large phylogenetic depth of the harsh-evolved populations, where the repeated bottlenecks drive the populations along a kind of forced march across the mutational landscape.

**Fig 16.**
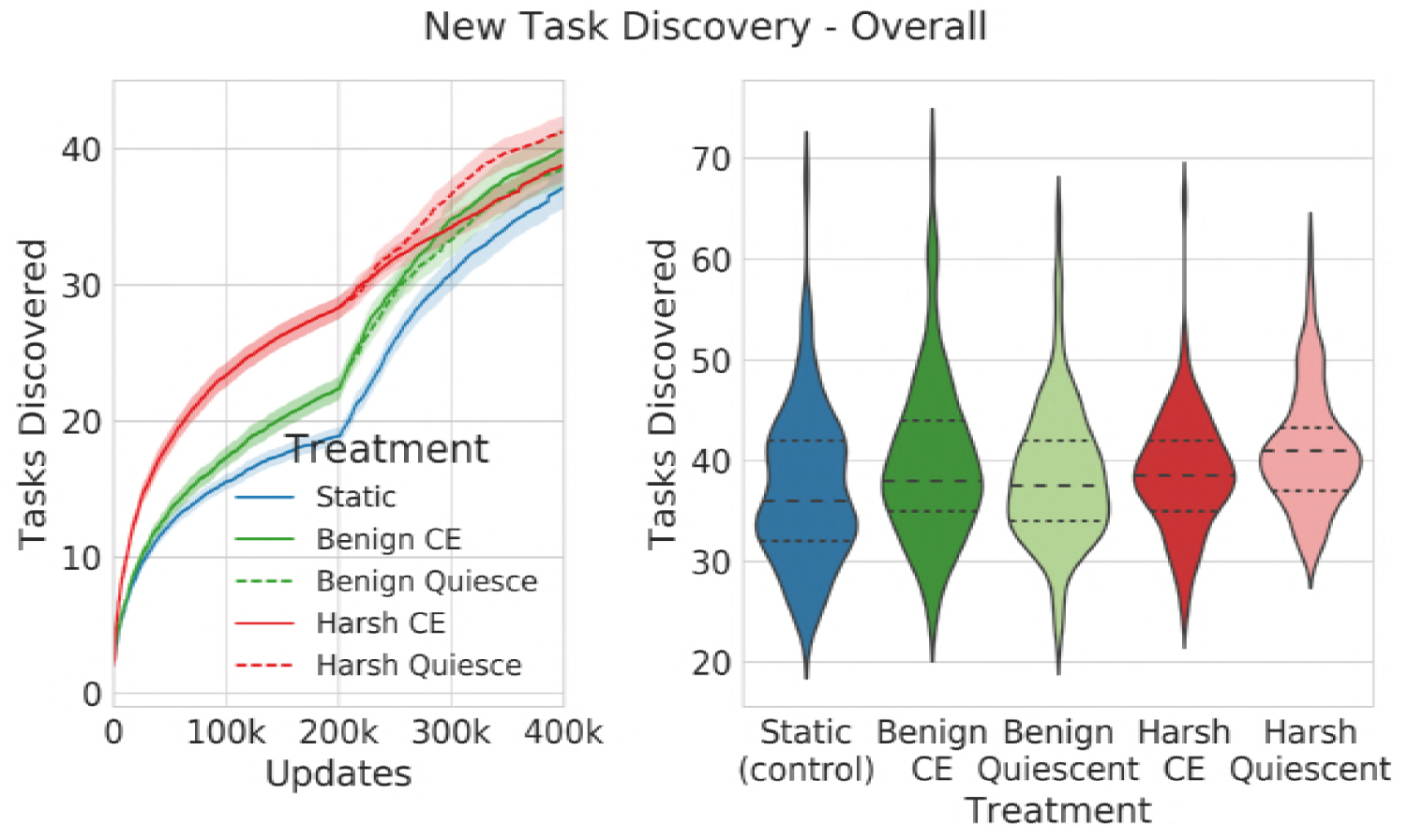
**Number of new expanded task set tasks discovered over the whole experiment**. The left plot shows a time-series of all new tasks discovered over the course of the entire run, including non-rewarded **expanded task-set** tasks. The right-hand plot shows the final count at the end of the run. Before we introduce rewards for performing the **expanded task-set** tasks, the harsh changing environment discovers far more new tasks (Mdn = 28.0, CI 95% [27.0, 30.0]) than either of the other treatments (Mdn = 22.0, CI 95% [22.0, 23.0]) (Wilcoxon Rank Sum Test: Z = 8.61, p << 0.001). These tasks appear despite no reward being given for performing any of the **expanded task-set** in the first part of the experiment.

However, as the experiment proceeds, and **expanded task-set** task rewards are introduced, this effect disappears, and task discovery rates converge (Kruskal-Wallis: H(2) = 6.97, p = 0.03).

### Task Performance

In addition to task discovery, task performance is another important measure of long-term evolvability, in that it quantifies exploitation and fixation of traits that are beneficial in new environments. We measured task performance in each of the changing environment treatments.

### Benign changing environments outperform harsh environments in task performance

Similar to task discovery, populations evolving in harsh changing environments performed far fewer distinct tasks than either the control, or either benign populations (Wilcoxon Rank Sum Test: Z = −11.22 and −11.15 respectively) (Fig 17). While both the BenignCE and BenignQuiescent populations seemed to outperform the control, the differences were not statistically significant (Kruskal-Wallis: H(2) = 2.76, p = 0.25).

**Fig 17.**
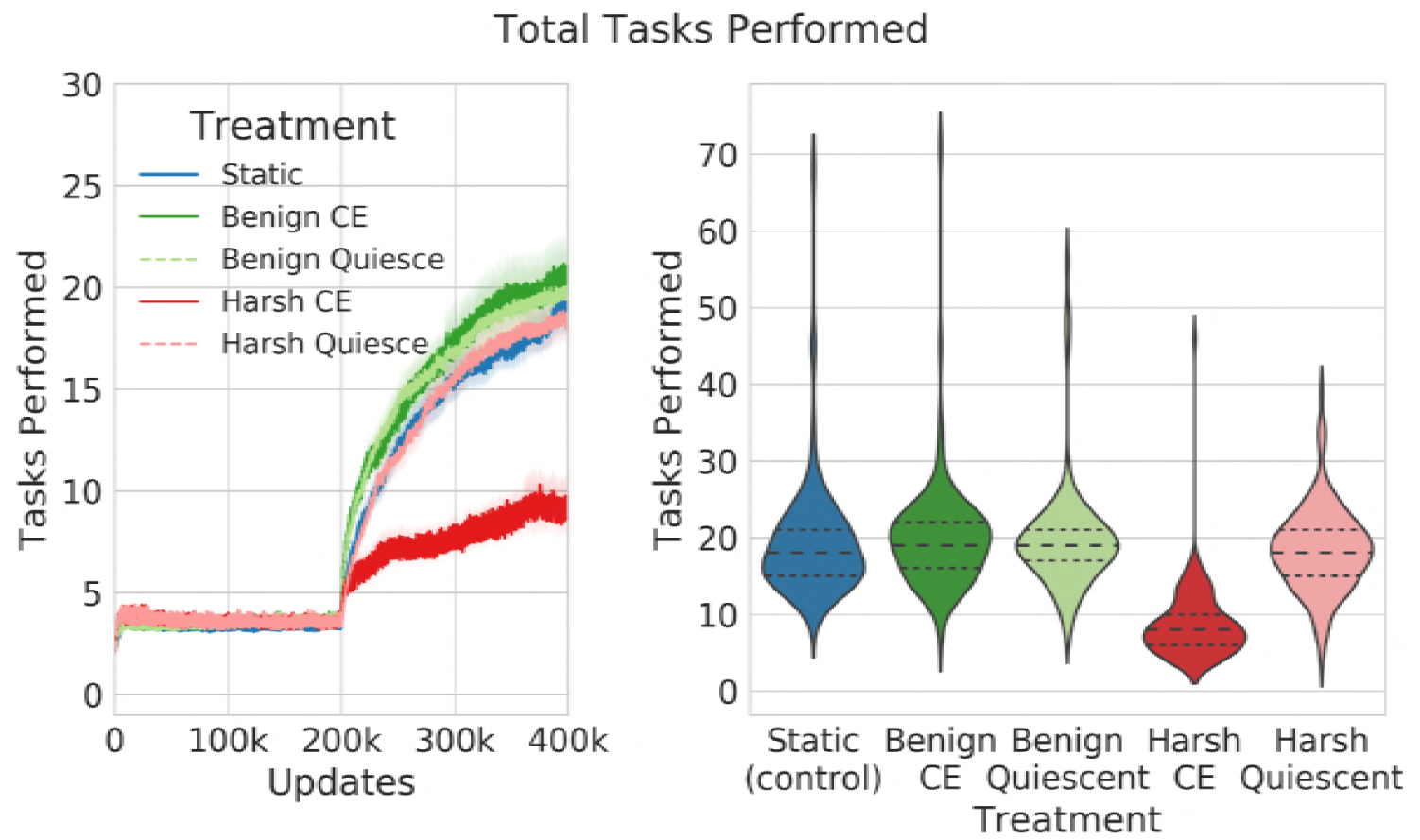
**Number of distinct tasks performed**. The left plot shows a time-series of the number of distinct tasks performed by the treatment populations over time. The right-had plot shows the number of tasks performed at the end of the experiments. The harsh changing environment treatment performs substantially and significantly fewer tasks than any of the benign or control treatments (Wilcoxon Rank Sum Test: Z = −11.22 and −11.15 respectively, p << 0.001). The benign treatments perform best, but the differences are not statistically significantly different than the control (Kruskal-Wallis: H(2) = 2.76, p = 0.25).

## Conclusion

In cyclic changing environments, the direction of selection shifts frequently, and periodically drives populations to not only explore new regions of the genetic landscape, but also to carry with them vestigial genetic information about previous environmental conditions. Thus, the resulting populations are not only adapted to the current environment, but also to the meta-environment of cyclic change. Because of their evolutionary history, the genomes contain vestigial fragments of genetic material that were adapted to prior environments. As this exploration proceeds, mutations accumulate in the population, each creating a link to a new region of the mutational landscape. As these links accumulate, they form a reservoir of mobility for the population to quickly shift to new phenotypes as dictated by current selective conditions. In this way, the accumulation of vestigial or pseudogene-like regions acts as an indirect adaptation to the larger pattern of changing selective forces.

By contrast, in static (non-changing) environments, the majority of neutral mutations do not connect to as many phenotypically-interesting regions of genotype-space. There are far fewer pseudogene-like regions available that could regain functionality should conditions change. Thus, populations evolved in static environments are less evolvable in the short-term.

These results suggest, therefore, that architectural features that help with short-term evolvability are more likely the result of repeated hitchhiking on adaptive mutants. In particular, we observed that much of the task-loss associated with the harsh changing environment could be attributed to increasing task length which is a result of the continuous addition of new mutations activating and deactivating the task. Despite this correlation, however, we observed a potential difference in robustness between the `XOR` and `EQU` tasks, which suggest that a kind of anti-robustness may also be selected for as a result of the changing environments.

### Long-Term Evolvability

The relationship between short- and long-term evolvability is non-obvious. Architectural features and selective pressures that promote repeated re-adaptation to a known set of environments may not be beneficial for the acquisition of entirely new adaptive traits, and the outcomes depend on the evolutionary and selective history of the population.

For example, harsh changing environments depress both fitness and population diversity, which might make these populations less effective at adaptation when introduced into a new environment. Even so, our results suggest that there are important architectural features conveyed by these environments that are beneficial for new task acquisition, despite the short-term downsides. Our experiments show that harsh changing environments, with their strong selective pressures, initially suppress the ability of populations to acquire new, weakly-selected traits. But if alternating selection is then removed, these populations are able to bounce back and rapidly acquire new tasks.

In contrast, in conditions where alternating selection persists, benign changing environments win at new task acquisition. Benign changing environments, with their milder set of selective pressures, are able to leverage their accumulated heritage of dormant vestigial sites to rapidly respond to selection, and acquire new tasks at a faster rate than either harsh or non-change-evolved populations.

### Limitations of Cyclic Changing Environments

Changing environments produce a set of selective pressures that speed up exploration of genotype space, while also building reservoirs of partial functionality that may be co-opted in the evolution of more complex structures. These features make changing environments useful for both their exploratory power in natural evolution, and as practical tools in the Artificial Life toolkit. Ultimately, however, as alluded to above, cyclic changing environments only re-tread existing phenotypic ground, and though genotypic exploration can be faster than under purely directional or stabilizing selection, the space explored remains constrained by the type of phenotypes that are selected. Despite this constraint, however, we see that, particularly under harsh conditions, a lot of novel genotypic ground may be explored, even without direct selection for novelty.

Even so, there must exist methods of exploring genotype space that do not suffer from these limitations at all. For example, perhaps repeated bottlenecking of populations could promote faster traversal of the fitness landscape in quasi-random directions. More ambitiously, perhaps these kinds of environments could be coupled with dynamically increasing open-ended complexity goals, or divergent selection mechanisms such as negative frequency dependence to promote the maintenance of diversity in evolving populations.

Understanding the mechanisms by which select environmental conditions alter fitness landscapes is vital to understanding the forces that promote evolvability and increase complexity. In particular, understanding the role of vestigial sites may help us untangle how robustness can promote evolvability. Are these vestigial sites merely inactive remnants, reservoirs of function, or are they part of a complex compensatory framework supporting and buffering the expression of the phenotype? Or all of these things? Changing environments provide one view into these dynamics, but we must explore further to find other mechanisms for exploring and exploiting genotype space.

## Acknowledgments

We would like to thank Alex Lalejini for helpful discussions and comments about the relationship between phenotypic plasticity and contingency loci, Joshua Nahum for discussions of experiments in changing environments, and Emily Dolson, Brian Goldman, and Anya Vostinar for their comments on early manuscript drafts. This material is based in part upon work supported by the National Science Foundation under Cooperative Agreement No. DBI-0939454 to CO and a Graduate Research Fellowship to RCK. Any opinions, findings, and conclusions or recommendations expressed in this material are those of the author(s) and do not necessarily reflect the views of the National Science Foundation.

## Supplemental Materials

### Sampled Nearby Mutational Landscape

As noted in equation 6, the mutants in the nearby mutational landscape include those that have more than one mutation. However, for completeness, we performed an exhaustive landscaping of the single-step mutational landscape, which, by definition, only includes mutants with a single mutation (see Figure 10). In order to verify that our results are indeed representative of the expected genomic and phenotypic diffusion rates, we sampled the mutants in the nearby mutational landscape using all naturally occurring mutations, including multiple mutations in a single mutant.

Our results (see figure 18) were virtually identical, showing that the sampling approach and the exhaustive landscaping produce qualitatively indistinguishable results.

**Fig 18.**
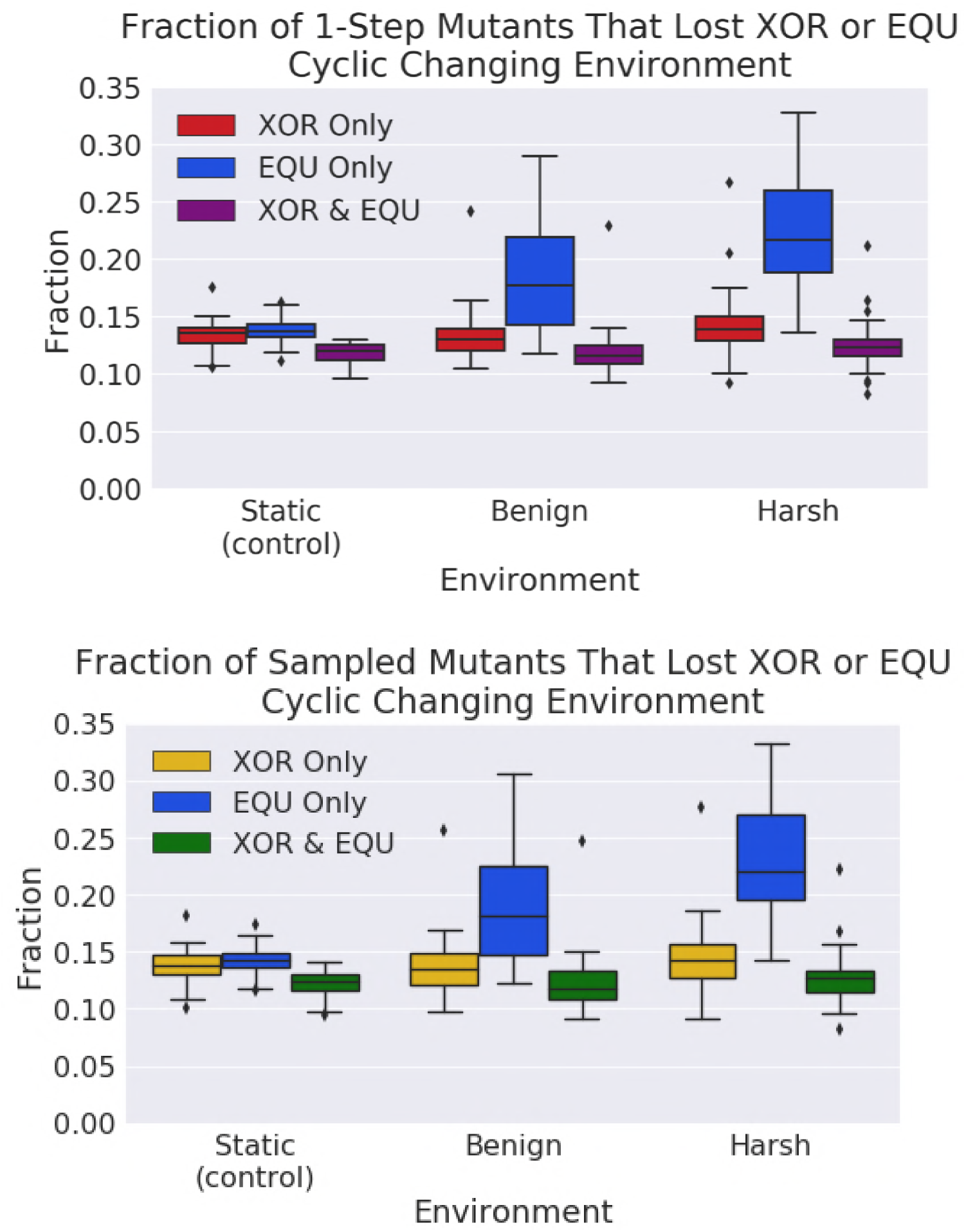
**A survey of the single-step and sampled mutational neighborhoods** around organisms that performed the fluctuating task. The results are qualitatively identical to each other.

## Footnotes

1 For simplicity, these measures are calculated based on the probability of single mutations occurring. However, in nature, multiple mutations may occur at once. We performed parallel experiments using a sampling approach (which could collect multiple mutations) to calculate similar metrics. These experiments yielded qualitatively similar results to those in this paper. See Section 1 of the supplemental materials.

2 As part of our initial controls, we hand-wrote an organism with separated sections that performed `XOR` and `EQU`. This hand-written organism had 121 instructions and, as such, we used this genome length as a constraint for the evolved organisms as well.

